# Biosynthesis of alkylcitric acids in *Aspergillus niger* involves both co-localized and unlinked genes

**DOI:** 10.1101/714071

**Authors:** Sylvester Palys, Thi Thanh My Pham, Adrian Tsang

**Author notes:** Contributed equally (Sylvester Palys, Thi Thanh My Pham).

## Abstract

Filamentous fungi are an abundant source of bioactive secondary metabolites (SMs). In many cases, the biosynthetic processes of SMs are not well understood. This work focuses on a group of SMs, the alkylcitric acids, each of which contains a saturated alkyl “tail” and a citrate-derived “head”. We initially identified their biosynthetic gene cluster and the transcriptional regulator (*akcR*) involved in the biosynthesis of alkylcitrates in the filamentous fungus *Aspergillus niger* by examining the functional annotation of SM gene clusters predicted from genomic data. We overexpressed the transcription regulator gene *akcR* and obtained from a litre of culture filtrate 8.5 grams of extract containing seven alkylcitric acids as determined by NMR. Hexylaconitic acid A comprised ~ 95% of the total production, and four of the seven identified alkylcitrates have not been reported previously. Analysis of orthologous alkylcitrate gene clusters in the Aspergilli revealed an in-cluster cis-aconitate decarboxylase gene (*cadA*) in *A. oryzae* and *A. flavus*, which in *A. niger* is located on a different chromosome. Overexpression of the *A. niger cadA* and *akcR* genes together shifted the profile of alkylcitrates production from primarily hexylaconitic acids to mainly hexylitaconic acids. We also detected two additional, previously unreported, alkylcitric acids in the double overexpression strain. This study shows that phylogenomic analysis together with experimental manipulations can be used to reconstruct a more complete biosynthetic pathway in generating a broader spectrum of alkylcitric compounds. The approach adopted here has the potential of elucidating the complexity of other SM biosynthetic pathways in fungi.

## Introduction

Filamentous fungi are a rich source of secondary metabolites (SMs). Many fungal SMs have useful bioactive properties; for example, the antibiotic penicillin, the cholesterol-lowering drug lovastatin, and the anti-cancer compound griseofluven^1–3^. In fungi, genes involved in the biosynthesis of SMs are typically co-localized in the genome and they are referred to as SM gene clusters^4^. These SM gene clusters generally consist of a “backbone” gene and multiple “tailoring” genes^5–6^. The backbone genes encode enzymes including polyketide synthases, non-ribosomal peptide synthetases, polyketide/nonribosomal peptide hybrid enzymes, dimethylallyl tryptophan synthases, terpene cyclases, and fatty acid synthases. The backbone enzymes generate the core of a particular set of SM compounds, serving as a scaffold for further modifications by tailoring enzymes^6^. Tailoring enzymes are thus responsible for most of the diversity of SMs^5^ and can serve as potential targets of manipulation for diverting secondary metabolism away from or towards particular compounds^7^. Secondary metabolite gene clusters may contain a transporter gene which facilitates the export of SMs out of the cell^8–10^. Lastly, a SM gene cluster can also contain gene(s) encoding transcriptional regulator(s) which may facilitate the transcription of the entire cluster and start the process of SM biosynthesis^11^. Overexpression of these clustered transcription factor genes can be a useful strategy to activate a SM gene cluster^12^.

Fungal orphan compounds are compounds that have been isolated from fungal cultures with no known genetic underpinnings. In *A. niger* and closely related black Aspergilli, approximately 140 orphan compounds have been isolated under laboratory conditions^13–14^. These orphan compounds can potentially be linked to their biosynthetic gene clusters by inferring the type of enzymes that are likely to be involved in their production, then locating a cluster(s) with the genes that code for those enzymes. This strategy has been successfully applied to locate the gene cluster of kotanin from *A. niger*^15^ and phomoidride from *Taleromyces stiptitatus*^16^. Tying orphan compounds to their SM gene clusters provides a method to study their biosynthesis, enhance their production, and manipulate the pathway toward desired products.

Multiple approaches have been devised over the past decades to identify and produce fungal SMs^17–18^. More recent approaches take advantage of the wealth of information garnered through whole genome sequencing and genome annotation to accelerate novel compound discovery^12, 15, 19–20^. The accumulated body of genomic information from many different organisms has allowed for the bioinformatic prediction of SM gene clusters. The predicted gene clusters have revealed that many organisms have far more potential regarding SM production than what has already been discovered. For example, only 12% of the curated SM gene clusters of *A. niger* have experimental support^13, 21^. In *A. nidulans* and *A. oryzae*, 13% and 3% of the predicted SM genes have experimental support respectively^13, 21^.

Alkylcitrates comprise two moieties; a saturated alkyl “tail” and a “head” derived from citric acid. Alkylcitrates isolated from filamentous fungi include the tensyuic acids (A-F)^22–23^, hexylaconitic acid anhydride^23–24^, hexylitaconic acid^24–25^, hexylcitric acid^26^ and hydroxylated hexylitaconic acids^27^ (Figure 1). Members of alkylcitric acids have been shown to possess useful bioactive properties including plant root growth promotion (hexylitaconic acid)^24^, anti-fungal (hexylaconitic acid anhydride)^28^, antibiotic, and anti-parasitic properties (tensyuic acid C)^22^. The production level of these bioactive alkylcitrates under laboratory conditions is low (ng-mg/L range) ^22, 24, 26–27^. In this work, we sought to use genomic information and chemical structure data to determine the SM gene cluster responsible for the production of the alkylcitric acid orphan compounds, manipulate their expression to increase the production of specific compounds, and to produce previously uncharacterized alkylcitric acids.

**Figure 1.**
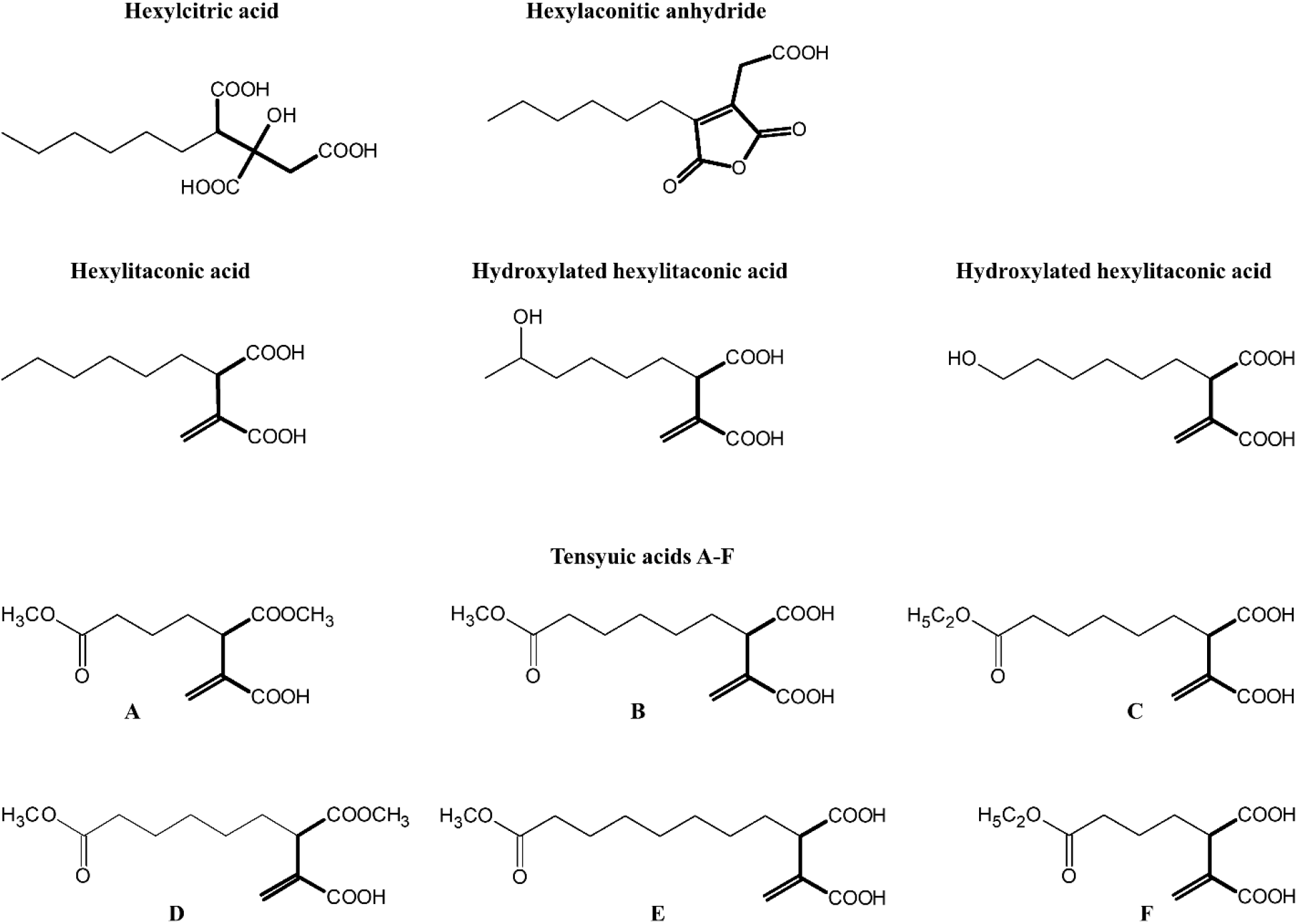
Structures of alkylcitric acids isolated from *A. niger*^22–24, 26–27^ showing two common moieties: a saturated hydrocarbon tail (thin bonds) and a citrate-derived head (thick bonds).

## Materials and Methods

### Alkylcitric acid gene cluster assignment and orthologous cluster analysis

The alkycitric acid biosynthetic gene cluster was assigned based on the annotation of secondary metabolite (SM) gene clusters in the *A. niger* NRRL3 genome available at the Genozymes website (www.fungalgenomics.ca). Orthologous alkylcitric acid clusters were located using the published Aspergilli genomes available at the Joint Genome Institute (JGI) MycoCosm website (https://genome.jgi.doe.gov/programs/fungi/index.jsf). BLASTP queries were carried out to locate orthologous gene clusters using the sequences of the fatty acid synthase backbone enzymes (gene IDs NRRL3_11763 and NRRL3_11767) and the citrate synthase enzyme (NRRL3_11764) from the predicted *A. niger* NRRL3 alkylcitric acid cluster. Clusters were considered orthologous to the *A. niger* NRRL3 alkylcitric acid gene cluster when the following three criteria were met: 1) protein sequence identity based on BLASTP of individual encoding genes was >50% and query coverage was >50%; 2) the orthologous genes are co-localized; and 3) the orthologous gene cluster contained fatty acid synthase backbone genes as well as citrate synthase and citrate dehydratase genes.

### Strains, culture conditions, and transformation protocols

The *A. niger* strain PY11 (N593 *glaA::hisG*)^29^ (Table 1) was used for production of alkylcitric acids. The DH5α strain of *Escherichia coli* was used for the propagation of cloned plasmids. Fungal cultures were initiated by inoculating spores (final concentration of 2×10^6^ spores/mL) into liquid minimal medium “J” (MMJ)^30^ in 96-well microplate at 250 µL per well or in petri dishes (14 centimeter in diameter (Sarstedt), containing 120 mL per plate). For growth of *A. niger* strains lacking a *pyrG* selection marker gene, MMJ media were supplemented with 10 mM uridine. For production of SMs, the cultures were incubated without shaking at 30°C for 5 days.

**Table 1.**
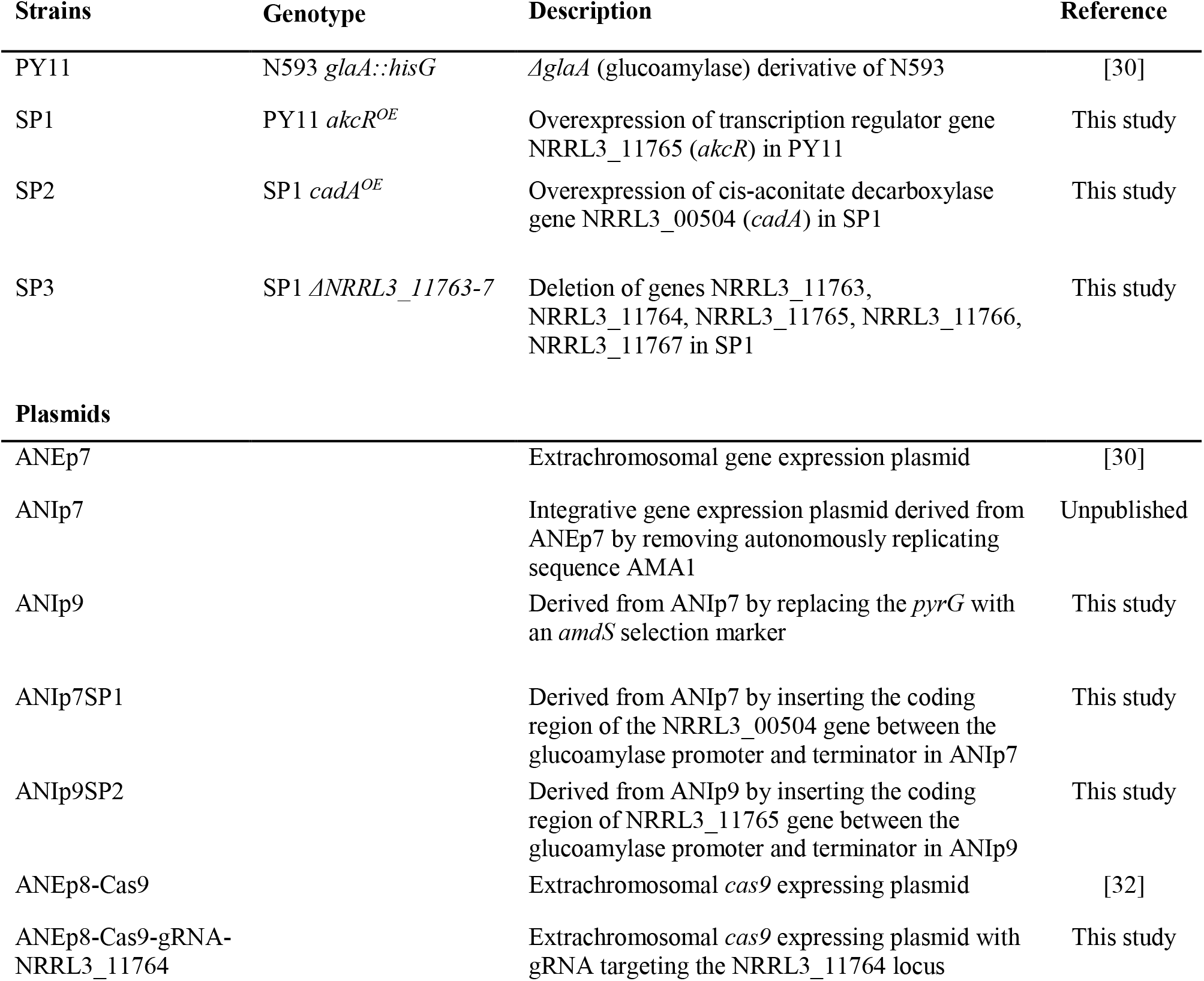
Strains and plasmids used in the study

Gene transformation in *E.coli* was carried out by heat shock of competent cells^30^. For transformation of *A. niger*, spores were inoculated into liquid complete media^31^ and shaken at 250 rpm for 18 hours prior to protoplasting. Protoplasting and transformation of *A. niger* were carried out as described^30^.

### Construction of overexpression cassettes for NRRL3_11765 and NRRL3_00504

The plasmids used in this study are listed in Table 1 while the primers used for PCR amplifications are listed in Table S1. The plasmid ANIp9 was used to construct the vector for the overexpression of the fungal-specific transcription factor gene NRRL3_11765. To amplify NRRL3_11765, primers PR_1 and PR_2 that have gene-specific sequences and an additional 22 nucleotides of adapter sequence were used. The NRRL3_11765 gene was amplified from genomic DNA isolated from *A. niger* strain N593^32^, and purified with a GeneJET Genomic DNA Purification Kit (Thermo K0721, Thermo Scientific, Grand Island, NY USA). The ANIp9 plasmid was amplified by PCR using primers Pr_3 and Pr_4 that contain an additional 22 nucleotides which are complementary to the adapter sequence of Pr_1 and Pr_2. The amplified vector and insert fragments were annealed and transformed into *E. coli* for propagation^33^. The resulting plasmid was transformed into the *A. niger* strain PY11, creating strain SP1 that overexpresses NRRL3_11765. Positive colonies were screened based on their ability to grow on acetamide as the sole source of nitrogen^34^.

The plasmid ANIp7 was used to construct the vector for the overexpression of NRRL3_00504, employing the same approach as for the NRRL3_11765 overexpression vector. The NRRL3_00504 gene was amplified using primers Pr_5 and Pr_6 while ANIp7 was amplified with Pr_3 and Pr_4. The resulting overexpression vector was transformed into strain SP1, creating strain SP2 which overexpresses NRRL3_00504 in the NRRL3_11765^OE^ background. Positive colonies showed growth on transformation plates containing no uridine.

### Construction of a deletion cassette for the removal of genes NRRL3_11763-7

A linear DNA cassette was designed to delete five genes encompassing NRRL3_11763 – NRRL3_11767 in the predicted alkylcitrate biosynthetic gene cluster. The deletion cassette contained 1255 base pairs (bp) of flanking DNA upstream of NRRL3_11763 (5’ flank) and 1331 bp of DNA downstream of NRRL3_11767 (3’ flank). Primers PR_7 and PR_8 were used to amplify the 5’ flank and primers PR_9 and PR_10 were used to amplify the 3’ flank. The 5’ and 3’ flanks were joined by overlap PCR^35^ using 22 nucleotide adapter sequences from primers PR_8 and PR_9 which are complementary to each other (Figure S1). The deletion cassette was co-transformed into strain SP1 (NRRL3_11765^OE^) along with the CRISPR/Cas9 vector ANEp8-Cas9-gRNA-NRRL3_11764. The CRISPR/Cas9 system was co-transformed to induce a double stranded break in the NRRL3_11764 gene and facilitate homologous recombination of the deletion cassette^31^. Construction of the gRNA insert for the CRISPR/Cas9 plasmid was carried out using primers PR_11 and PR_12 as previously described^31^. Screening for the deletion was done by multiplex PCR with primers Pr_13 and Pr_14 that bind to the genome outside the 5’ and 3’ homologous regions, respectively, and primers Pr_15 and Pr_16 that bind to the NRRL3_11765 gene inside the deletion (Figure S1). The expected band for the deletion strain is 2697 bp with primers Pr_13 and Pr_14, and no amplification of the NRRL3_11765 gene (Pr_15 and Pr_16). In the parental strain, the expected band for Pr_15 and Pr_16 is 1653 bp while for Pr_13 and Pr_14 no band is expected since the size of the amplicon is over 25 kb (Figure S1).

### Sample preparation and data analysis by liquid chromatography mass spectrometry

Following growth of the transformants and parental strain, 75 µL of growth cultures were collected in 1.5 mL microfuge tubes and centrifuged at 16,000 x g for 45 minutes to remove mycelia, spores and cellular debris. The supernates were transferred to new tubes and an equal volume of cold methanol (−20°C) was added for protein precipitation. Following incubation on ice for 10 minutes, samples were centrifuged at 16,000 x g for 45 minutes to remove precipitated proteins. Supernates were then transferred to fresh tubes and an equal volume of 0.1% formic acid was added.

Electrospray liquid chromatrography mass spectrometry (LC-MS) analyses were performed on a 7-Tesla Finnigan LTQ-FT-ICR mass spectrometer (Thermo Electron Corporation, San Jose, CA). Ionization voltage used was 4900 V in positive mode and 3700 V in negative mode. Scan range was from 100 to 1400 m/z at a 50000-resolution setting. The solvent delivery system used was a Series 200 auto sampler and micropump (Perkin Elmer, Waltham, MA). Injection volume was 10 µL and flow rate was 250 µL/minute. Reversed-phase liquid chromatography (RPLC) separation was performed using an Eclipse C18 3.5 µm, 2.1 x 150 mm column (Agilent, Santa Clara, CA). The solvents used to generate the gradient during RPLC separation were 0.1% formic acid in water for solvent A and 0.1% formic acid in acetonitrile for solvent B. The gradient was used to elute the metabolites: 5% B isocratic for 1 minute, increased to 95% B in 10 minutes, isocratic at 95% B for 1 minute, decreased to 5% B in 0.1 minute, and isocratic at 5% B for 5.9 minutes.

### Sample preparation and data analysis by gas chromatography mass spectrometry

For gas chromatography mass spectrometry (GC-MS) analysis, 0.1 mL of growth culture was extracted twice with 1 mL of ethylacetate. The extract was dried under a stream of nitrogen. Dried extracts were then dissolved in 50 µL of N,O-bis(trimethylsilyl)trifluoroacetamide (BSTFA) (Supelco, Sigma-Aldrich), followed by heating at 70°C for 30 min^36^. The trimethysilyl derivatized product in BSTFA was then injected in a Hewlett Packard HP6890 gas chromatograph equipped with a HP5975 mass detector and a DB-5MS column (25 m×0.2 mm) (Agilent Technologies). The program started at 80°C (held for 2 min), followed by an increase of 15°C per min to 315°C and held at 315°C for 3 min. The mass spectrometer was operated in the electron impact ionization mode.

### Secondary metabolite purification and structural analysis by nuclear magnetic resonance spectra

One litre of the growth cultures of strain SP1 (NRRL3_11765^OE^) were extracted with 1 L of ethylacetate followed by two additional extractions with 500 mL of ethylacetate each. The extract was dried *in vacuo* to yield a brown syrup. A portion of the material (850 mg) was dissolved in 2 mL of a 4:1 (v/v) mixture of acetonitrile and high performance liquid chromatography (HPLC)-grade water. The sample was injected (100 µL/injection) into an Atlantis dc18 OBD Prep Column (100Å, 5 µm, and 19 mm X 100 mm) (Waters) connected with an Agilent 1100 series HPLC at a flow rate of 20 mL/minute. Solvent A was HPLC-grade water containing 0.1% formic acid and solvent B was acetonitrile. The HPLC gradient included a linear increase from 5% to 95% of solvent B over the course of 10 minutes, and then remain at 95% of B for 3 minutes. Solvent B was returned to 5% within 1 minute and then held for 5 minutes to allow for column re-equilibration. The detector was set at a wavelength of 210 nm. Fractions were collected based on peak signals. The collected fractions were re-extracted with ethylacetate and dried to yield brown syrups ready for nuclear magnetic resonance (NMR) analysis.

Secondary metabolites from the strain SP2 (SP1, NRRL3_00504^OE^) were extracted and purified following the identical method used for strain SP1. These alkylcitric acids were eluted and collected from HPLC at different retention times.

For NMR analysis, the dried syrup materials were dissolved in deuterated chloroform or deuterated methanol (Table S2). All NMR spectra were recorded with a Varian VNMRS-500 MHz) at 25°C.

## Results

### Locating the biosynthetic gene cluster for the alkylcitric acids

To locate the gene cluster underpinning alkylcitric acid production, we began by examining the structures of the alkylcitric acids that have previously been reported (Figure 1). Their structures consist of a saturated hydrocarbon “tail” and a “head” derived from citric acid, suggesting that a fatty acid synthase or a highly reducing polyketide synthase is involved in the synthesis of the hydrocarbon tail while a citrate generating or modifying enzyme is involved in the production of the citrate moiety. Using the secondary metabolite gene clusters annotated in the genome of *A. niger* NRRL3, we located a single candidate gene cluster spanning genes NRRL3_11759-NRRL3_11769 (Figure 2). The cluster contains genes encoding a dienelactone hydrolase (NRRL3_11759), two fungal-specific transcription factor (NRRL3_11760 and NRRL3_11765), transporter (NRRL3_11761), fatty acid synthase alpha and beta subunits (NRRL3_11763 and NRRL3_11767), a citrate synthase (NRRL3_11764), a 2-methylcitrate dehydratase (NRRL3_11766), a polyprenyl synthase (NRRL3_11768) and an aldehyde dehydrogenase (NRRL3_11769).

**Figure 2.**
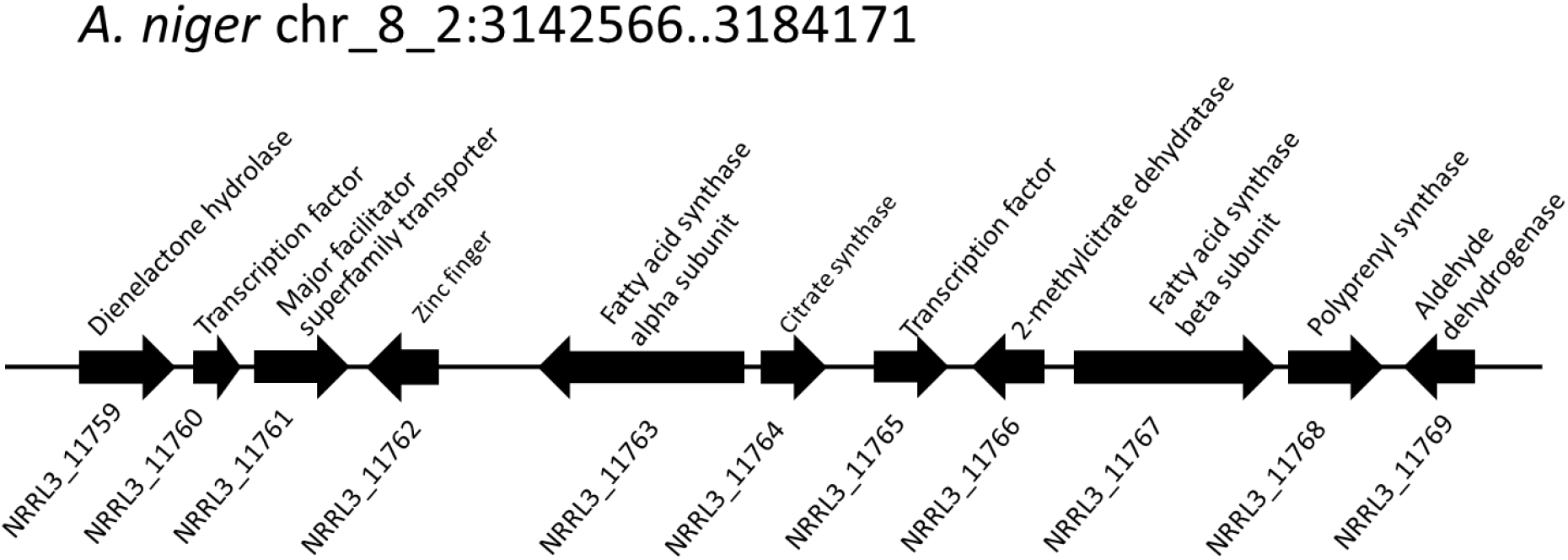
Predicted alkylcitric acid gene cluster of *A. niger*.

### Overexpression of transcription factor gene NRRL3_11765 leads to the overproduction of secondary metabolites

The gene NRRL3_11765 encoding a fungal-specific transcription factor is located in the predicted alkylcitric acid gene cluster, and is presumed to be the regulator of the alkycitrate biosynthesis genes. To overexpress NRRL3_11765, we placed the gene under the control of the maltose-inducible glucoamylase promoter^37^. The recombinant gene was transformed into *A. niger* for random integration into the genome. Following growth in the presence of the inducer maltose, we analyzed the extracellular medium of strain SP1 (NRRL3_11765^OE^) and the parental PY11 strain by GC-MS. Strain SP1 showed multiples peaks while extracellular medium of the parental strain did not show any of these peaks (Figure S2).

### Nuclear magnetic resonance (NMR) analysis reveals seven alkylcitric acids

For NMR analysis, extracellular culture fluid of strain SP1 was extracted with ethylacetate and dried. About 8.5 grams of oily material were obtained from 1 L of extracellular fluid. Secondary metabolites in the material were separated by HPLC and were eluted from 4.0 min to 10.0 min (Figure S3). To elucidate the structure of SMs, fractions from HPLC were subjected to NMR and mass spectrometry analyses.

We used one-dimensional and two-dimensional NMR spectra to elucidate the structure of the two major metabolites produced by strain SP1: hexylaconitic acid and hexylitaconic acid (Supplemental information, Section on structural elucidation). Structures of the other compounds were determined by comparing their ^1^H- and ^13^C-NMR spectra with the major hexylaconitic acid and hexylitaconic acid, and with published data^24, 26–27, 38^. Since all compounds in this study are derivatives of either hexylaconitic acid or hexylitaconic acid, we designate the major compounds as hexylaconitic acid A and hexylitaconic acid A. Their derivatives are then denoted with successive letters alphabetically. Seven compounds were identified from growth media of strain SP1 (NRRL3_11765^OE^), including hexylaconitic acids A and B, and hexylitaconic acids A to E. Compound structures are shown in Figure 3. Four of the seven compounds (hexylaconitic acids A, B and hexylitaconic acids A, E) have not been reported before. The major compound hexylaconitic acid A comprised about 95% of the crude extract, with the remaining 5% for all the other compounds combined. The NMR data of all seven alkylcitric acids are presented in Table S2. Since the seven compounds overproduced in strain SP1 (NRRL3_11765^OE^) are all alkylcitric acids, we conclude that NRRL3_11765 encodes the regulator of alkylcitric acids biosynthesis, and call this gene *akcR*.

**Figure 3.**
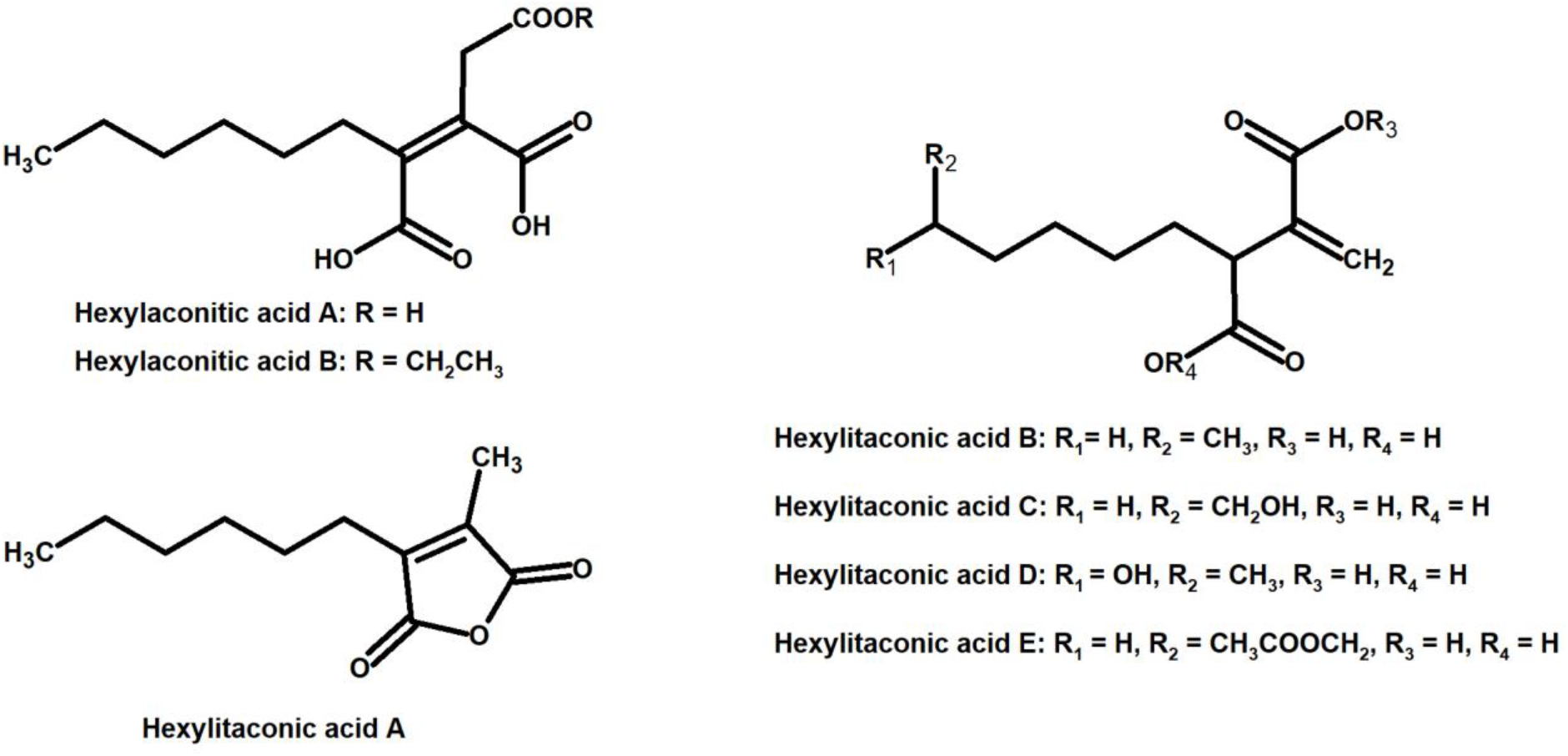
Compounds identified from growth media of strain SP1 (NRRL3_11765^OE^)

### Deletion of genes NRRL3_11763 to NRRL3_11767 abolishes the production of alkylcitric acids

To confirm the involvement of the predicted alkylcitric acid cluster in the production of alkylcitric acids, we designed a deletion cassette to remove five co-localized genes from the cluster, NRRL3_11763 to NRRL3_11767. These genes code for fatty acid synthase subunits A and B (NRRL3_11763 and NRRL3_11767), citrate synthase (NRRL3_11764), the transcription regulator *akcR* (NRRL3_11765), and 2-methylcitrate dehydratase (NRRL3_11766). The deletion was constructed in strain SP1 (*akcR*^OE^) background where the *akcR* overexpression allele is randomly integrated in the genome and remains active. Mass spectra of the cluster deletion strain did not detect peaks corresponding to any of the alkylcitric acids (Figure S4). Given the lack of alkylcitric acid production in strain SP3 (*akcR^OE^* ΔNRRL3_11763-7), we refer to this region of the genome as the alkylcitric acid gene cluster.

### Analysis of orthologous alkylcitric acid gene clusters reveals the involvement of cis-aconitate decarboxylase

Over 95% of the alkylcitric acids produced by strain SP1 were hexylaconitic acids and less than 5% were hexylitaconic acids (Figure S3). In the itaconic acid biosynthesis pathway, cis-aconitate decarboxylate converts aconitic acid to produce itaconic acid^39–40^. Hence we posited that the conversion of hexylaconitic acid to hexylitaconic acid requires an enzyme with cis-aconitate decarboxylase activity. To identify the candidate cis-aconitate decarboxylase, we examined orthologous gene clusters predicted to be involved in alkylcitric acids biosynthesis. Previous reports have shown that alkylcitric acids and the similar structured maleidrides are being produced not only in *A. niger* but also in other filamentous fungi including *Penicillium striatosporum*^27, 41^ and *A. oryzae*^42^. Therefore, we used the alkylcitric acid cluster of *A. niger* as query to search for orthologous gene clusters in the published Aspergilli genomes. Twenty orthologous alkylcitric acid gene clusters are found in section Nigri and two in section Flavi (*A. flavus* and *A. oryzae*) of the Aspergilli (Figure S5). The orthologous clusters are not identical (Figure 4). They share five genes in common, which we call: *akcA*, fatty acid synthase alpha subunit (NRRL3_11763); *akcB*, citrate synthase (NRRL3_11764); *akcR*, transcription regulator (NRRL3_11765); *akcC*, 2-methylcitrate dehydratase (NRRL3_11766); and *akcD*, fatty acid synthase beta subunit (NRRL3_11767).

**Figure 4.**
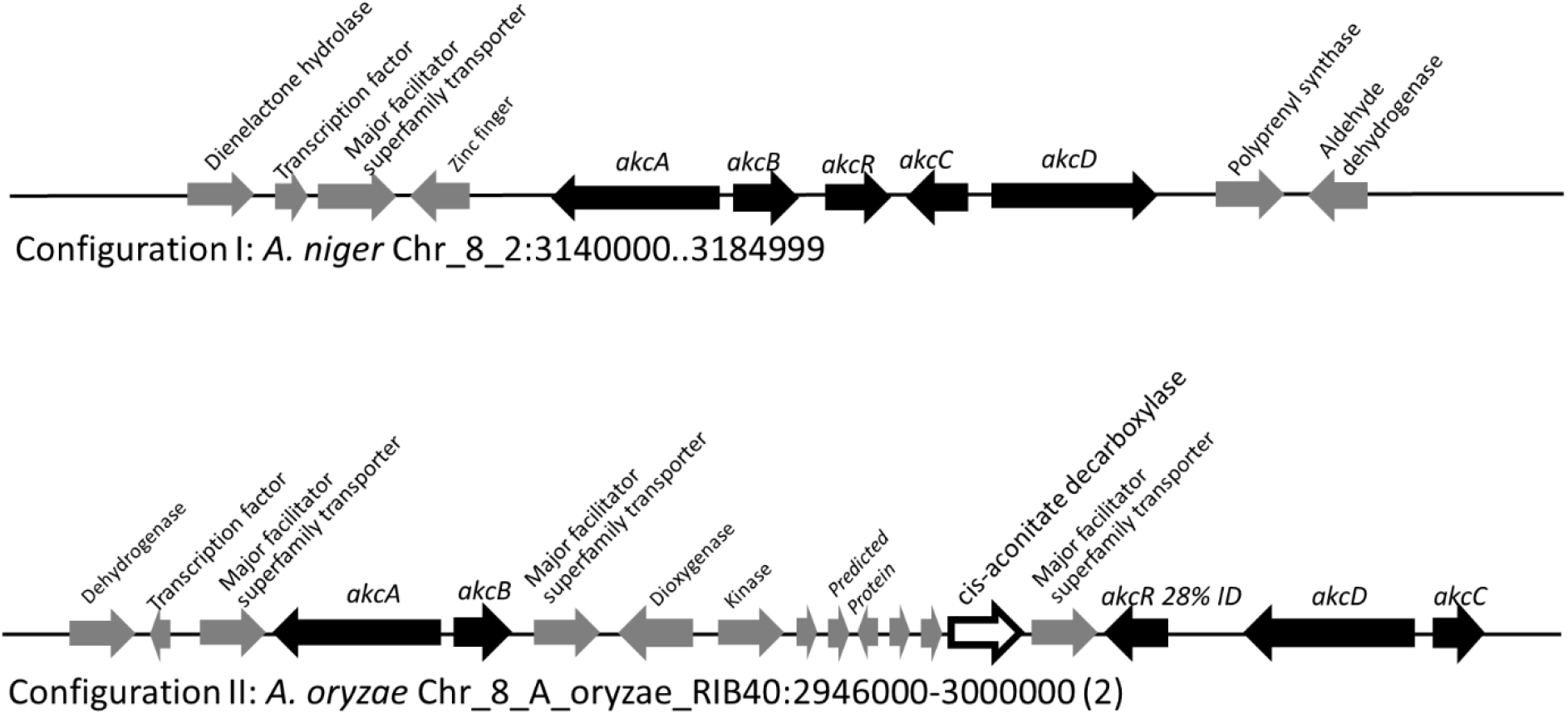
Two general configurations of the alkylcitric acid cluster in the Aspergilli. The percent identity (28%) displayed between *akcR* and the in-cluster transcription factor in *A. oryzae* represents a BLASTP top hit for *akcR*, but which is below the cut off (% identity >50%) to be considered orthologous.

There were two configurations for the predicted alkylcitric acid gene clusters (Figure 4). The first is found in section Nigri, as exemplified in *A. niger* NRRL3, contains the five conserved genes (conserved cluster) in a contiguous arrangement. The second is found in section Flavi, as exemplified in *A. oryzae*, there are intervening genes within the conserved clusters.

Notably, a gene predicted to encode cis-aconitate decarboxylase is in the alkylcitrate cluster of the genomes of the *A. flavus* and *A. oryzae*. The *A. niger* orthologue of this cis-aconitate decarboxylase is NRRL3_00504 on Chromosome I, outside the alkylcitric acid cluster which is on Chromosome VIII. This gene may potentially be involved in alkylcitric acid biosynthesis.

### Overexpression of NRRL3_00504 gene in *akcR^OE^* strain shifts production from hexylaconitic acid A to hexylitaconic acids

In the SP1 (*akcR*^*OE*^) strain, hexylaconitic acid A constitute about 95% of total alkylcitric acids with its downstream products including hexylitaconic acids comprising less than 5%. This observation suggests that the abundance of hexylaconitic acid produced by the SP1 (*akcR*^*OE*^) strain is the result of a biosynthetic bottleneck. As in the itaconic acid biosynthesis pathway^39–40^, we hypothesized that cis-aconitate decarboxylase mediates the conversion of hexylaconitic acids to hexylitaconic acids. The gene NRRL3_00504 is predicted to encode cis-aconitate decarboxylase and it is the closest orthologue to the predicted cis-aconitate decarboxylase genes found in the alkylcitrate gene clusters of *A. oryzae* and *A. flavus*. To examine the involvement of NRRL3_00504 in the biosynthesis of hexylitaconic acids, we overexpressed NRRL3_00504 in strain SP1 (*akcR*^*OE*^). As in the case of strain SP1, we obtained about 8.5 grams/L of metabolites in strain SP2 (*akcR*^*OE*^ NRRL3_00504^OE^). However, in strain SP2 about 10% of the metabolites are hexylaconitic acid A and 65% of the metabolites are hexylitacinic acids (B, C and D) (Figure 5). Since the shift of production of hexylaconitic acids to hexylitaconic acid is mediated by NRRL3_00504, the result provides experimental evidence that NRRL3_00504 encodes cis-aconitate decarboxylase. We therefore rename NRRL3_00504 as *cadA*.

**Figure 5.**
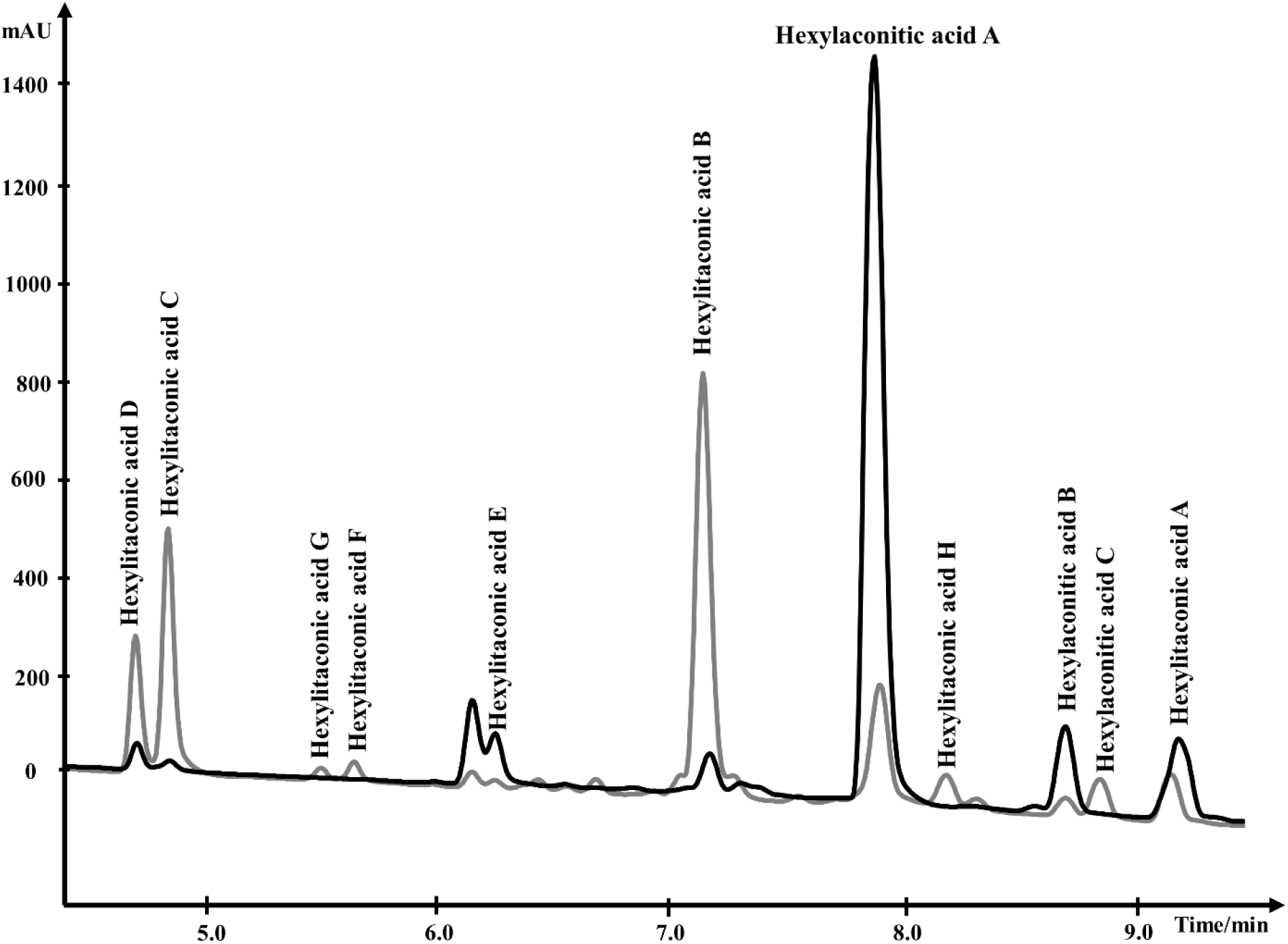
Overlapping HPLC chromatograms of products produced by strain SP1 (*alkR*^OE^), in black, and by strain SP2 (*akcR*^*OE*^*cadA*^*OE*^), in gray.

Further, strain SP2 (*akcR*^*OE*^*cadA*^*OE*^) produces four additional alkylcitric acids: hexylaconitic acid C and hexylitaconic acids F to H (Figure 5 and 6). Two of four compounds (hexylaconitic acid C and hexylitaconic acid F) have not been reported before. Each of these compounds was produced at ~0.2-0.3 g/L range (Figure 7). The NMR data for these compounds are shown in Table S2.

**Figure 6.**
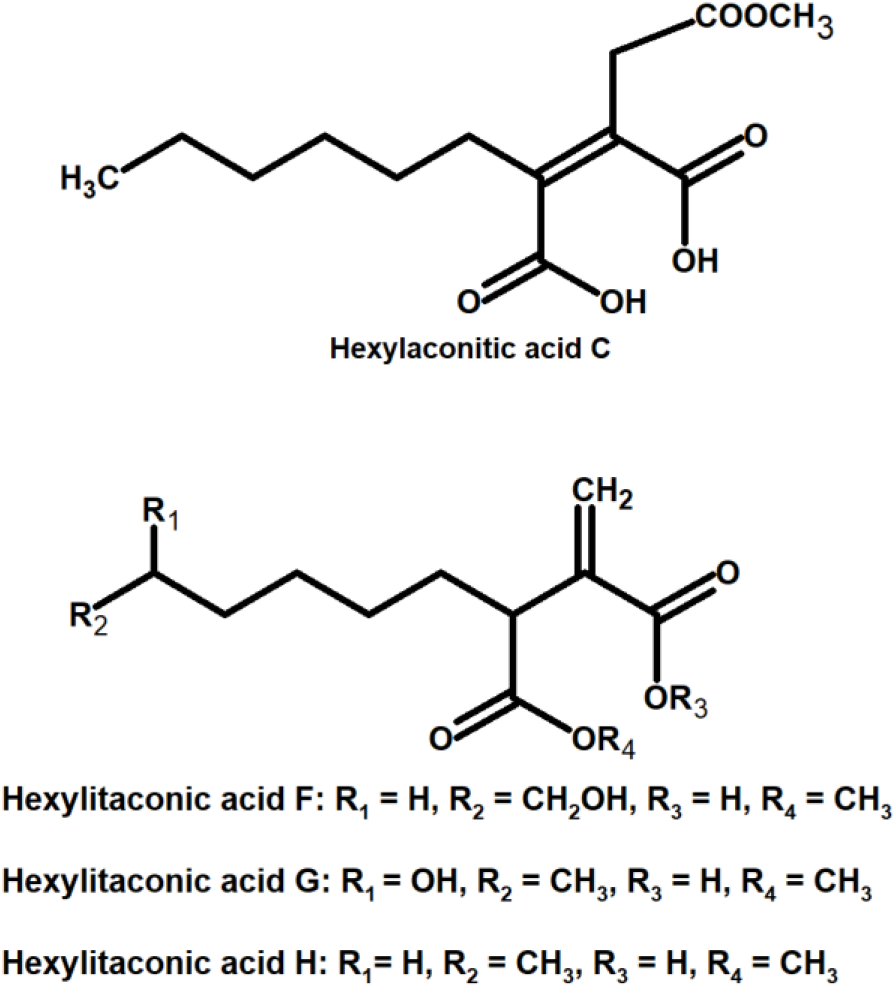
Compounds identified from growth media of strain SP2 (*akcR*^*OE*^ *cadA*^*OE*^)

**Figure 7.**
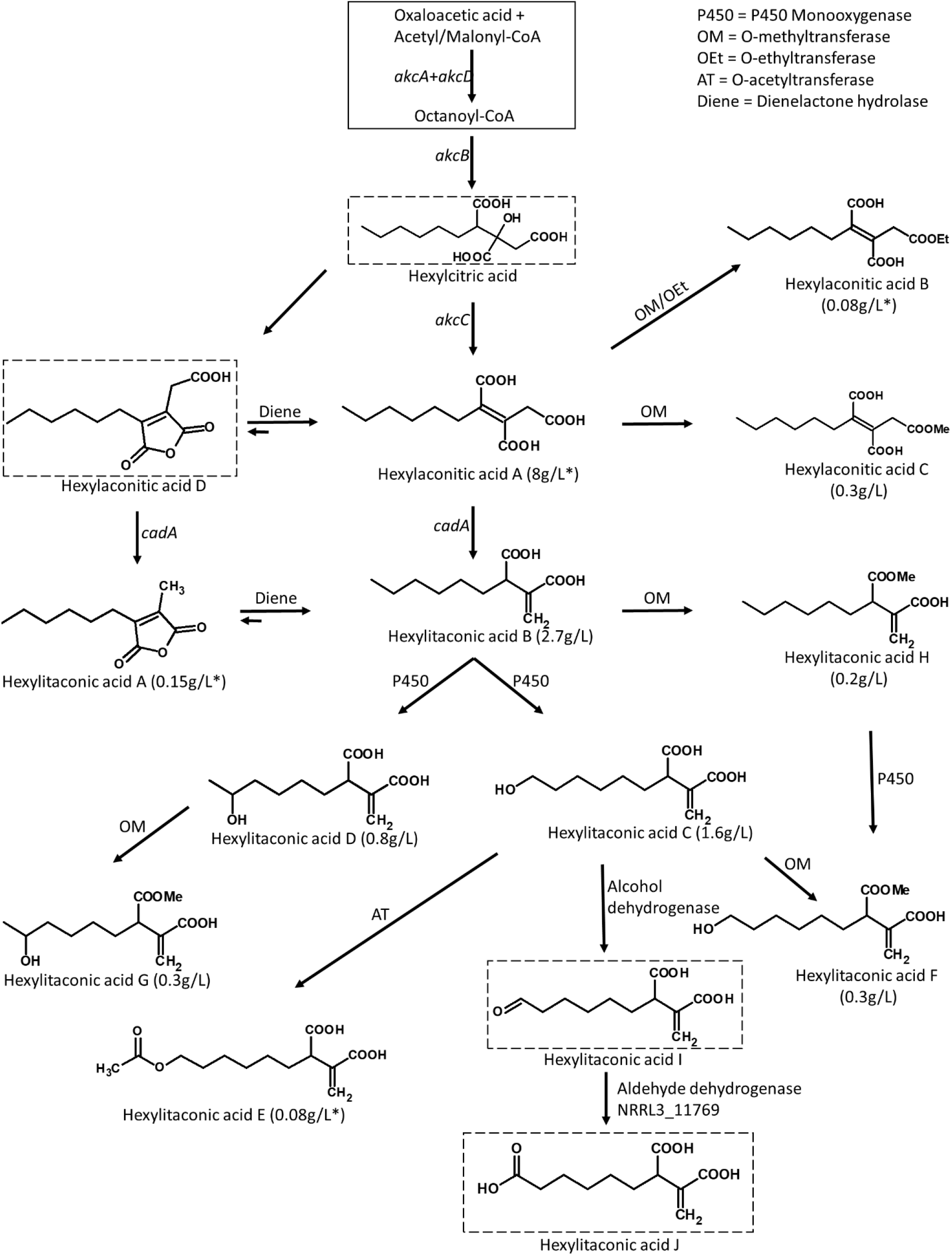
Predicted biosynthetic pathway for the alkylcitric acids. The solid box represents building blocks of hexylcitric acid, which were not detected in this work. Dashed boxes indicate compounds which were detected by MS; all other compounds were purified and identified by NMR. Yields indicated with an asterisk (*) represent values which were isolated from the SP1 (*akcR*^*OE*^) strain; all other values represent yields which were obtained from the SP2 (*akcR^OE^ cadA^OE^*) strain.

## Discussion

Overexpression of the transcription factor *akcR* and the cis-aconitate decarboxylase gene *cadA* led to the production of 11 alkylcitric acids. Based on the chemical structure of the alkylcitric acids resolved by NMR and the genes co-localized in the genome of *A. niger* and related species in section Nigri, we hypothesize that the biosynthetic pathway for alkylcitric acids involves at least six co-localized genes and one unlinked gene: *akcA* and *akcD* (NRRL3_11763 and NRRL3_11767) for the two subunits of fatty acid synthase, *akcB* (NRRL3_11764) for citrate synthase, *akcC* (NRRL3_11766) for 2-methylcitrate dehydratase, NRRL3_11759 for dienelactone hydrolase, NRRL3_11769 for aldehyde dehydrogenase, and the unlinked *cadA* (NRRL3_00504) for cis-aconitate decarboxylase. The reconstructed pathway is shown in Figure 7.

The pathway is predicted to begin with the production of hexylcitric acid, which is generated by a citrate synthase (*akcB*) and fatty acid synthase (*akcA* and *akcD*). Despite finding peaks in LC-MS data ([M+H]^+^ = 277.1287) (Figure S7), we were unable to purify and confirm the structure of hexylcitric acid by NMR. This initial compound may be efficiently converted to downstream hexylaconitic acid A and only present in small quantities. The next step in the pathway is the dehydroxylation of hexylcitric acid by 2-methylcitrate dehydratase (*akcC*) to produce hexylaconitic acid A. Finally, a decarboxylation reaction occurs to yield hexylitaconic acid B. This step was carried out by cis-aconitate decarboxylase (*cadA*) whose role was confirmed in this study.

The alkylcitrate pathway shares similarity with the itaconic acid pathway from *A. terreus*^39–40^, since they share a similar chemical structure and gene annotations. The itaconic acid pathway starts by the production of citric acid from acetyl-CoA and oxaloacetic acid followed by dehydroxylation to aconitic acid and finally a decarboxylation to itaconic acid. These steps are carried out by a citrate synthase, a dehydratase and a cis-aconitate decarboxylase respectively^39–40^. The biosynthetic pathway of itaconic acid and the proposed pathways for hexylitaconic acid B are shown in Figure S8.

The next steps of the alkylcitrate biosynthetic pathway may involve an omega oxidation reaction. This three-step reaction^43^ would require one or more P450 monooxygenase(s) to generate tail end hydroxyl groups (hexylitaconic acids C and D). An alcohol dehydrogenase then oxidizes the hydroxyl group to generate an aldehyde (hexylitaconic acid I). Finally an aldehyde dehydrogenase is predicted to generate a carboxyl group (hexylitaconic acid J)^43^. Although we could not confirm the structures of hexylitaconic acid I and J by NMR, we detected their molecular masses ([M+H] ^+^ = 229.1076 and [M+H] ^+^ = 245.1025 respectively) by LC-MS (Figure S7). While we do not know the locations in the genome for the P450 monooxygenase(s) and the alcohol dehydrogenase, an aldehyde dehydrogenase (NRRL3_11769) is co-localized with the alkylcitric acid gene cluster (Figure 2). As in the case of *cadA*, the omega oxidation step and perhaps other biosynthetic reactions in the alkylcitric acid pathway appear to be generated by enzymes whose genes do not co-localize. We do note, however, that not all clustered genes have been completely ruled out as candidates for these steps.

Other genes which co-localize with the alkylcitric acid cluster may also be involved in alkylcitric acid biosynthesis. In particular, the clustered gene NRRL3_11759 encoding a dienelactone hydrolase, an anhydride ring-opening enzyme^44^, may be responsible for shifting the ringed hexylaconitic acid D and hexylitaconic acid A towards the non-ringed hexylaconitic acid A and hexylitaconic acid B (Figure 7). This would provide an explanation for why the non-ringed forms were more abundant than the ringed forms. For instance, the non-ringed form (hexylaconitic acid A) was detected as major compound in the SP1 (*akcR*^*OE*^) extract (~ 95% of total extract) while its ringed form (hexylaconitic acid D) was detected by LC-MS ([M+H] ^+^ = 241.1076) (Figure S7) but not by purification process, indicating a lower level of production. Similarly, hexylitaconic acid B was the major compound (~ 35% of total extract) of the SP2 (*akcR*^*OE*^ *cadA*^*OE*^) strain while its ringed form (hexylitaconic acid A) was isolated at ~ 1.5% of the total extract.

In addition to the tail modifications, we also detected one O-ethylated (hexylaconitic acid B), one O-acetylated (hexylitaconic acid E) and four O-methylated (hexylaconitic acid C and hexylitaconic acids F to H) alkylcitric acids indicating other enzymes capable of appending the hydroxyl and carboxyl groups of the alkylcitric acids may also be involved in the pathway. No such transferase genes are located nearby or inside the alkylcitric acid cluster and we are also unable to locate them in orthologous clusters in other filamentous fungal species.

Conversely, there are 6 previously published alkylcitric acids (tensyuic acids A-F) (Figure 1) that we did not detect from our strains (SP1 and SP2). Tensyuic acids B to D appear to be o-methylated and o-ethylated forms of hexylitaconic acid J. The low production of hexylitaconic acid J and its direct precursor (hexylitaconic acid I) would provide an explanation for the absence of tensyuic acid B to D in our samples, despite identifying other o-methylated and o-ethylated compounds of the pathway. Locating and overexpressing the responsible alcohol dehydrogenase and aldehyde dehydrogenase genes should therefore help to increase the production of tensyuic acid B to D. Tensyuic acids A and F are products of o-methylation and o-ethylation of the 4-carbon-tail itaconic acid while tensyuic acid E is an o-methylated form of the 8-carbon-tail itaconic acid. There is no published evidence supporting the ability of one fatty acid synthase to synthesize compounds containing different carbon chain lengths. We therefore predicted that tensyuic acids A, E and F were synthesized by different fatty acid synthases, which are active in the *A. niger* FKI-2342 strain (tensyiuc acid producer^22^), but not in our strains (SP1 and SP2).

The common conception of fungal secondary metabolism has been that natural products are generated from discrete clusters of genes^45–47^. As our understanding increases we begin to see more exceptions where SM genes are not co-localized. For example, the genes underpinning melanin production are clustered in *A. fumigatus*, while in *A. niger* the genes are scattered throughout the genome^48^. The involvement of both co-localized and unlinked genes in SM biosynthesis has implications for heterologous expression as well as pathway reconstruction. Experimental design that focuses solely on clustered genes^49–53^ may fail to realize the full diversity of SM generation or result in partial pathway reconstructions. Analyzing orthologous clusters can help address some of these shortcomings by identifying unlinked genes that are involved in SM biosynthesis.

The overexpression strains SP1 (*akcR*^*OE*^) and SP2 (*akcR*^*OE*^ *cadA*^*OE*^) generated alkylcitric acids at the g/L level (defined here as artificial production) compared favourably to the levels of production previously reported from *A. niger* and other species (isolated without genetic manipulation and defined here as natural production)^54^. For example, natural production of hexylitaconic acid B yielded 0.5 mg/L in *Penicillium striatisporum*^27^, 14 mg/L in *A. niger* K88^25^ and 14 mg/L in *A. niger* AN27^24^. In our study, artificial production of hexylitaconic acid B yielded 214 mg/L in strain SP1 and increased to 2.7 g/L in SP2. This was 200 fold higher than that from natural production in other strains of *A. niger* and more than 5000 fold higher than *P. striatisporum* under natural production. Hexylaconitic acid A was produced at 8 g/L by the SP1 (*akcR*^*OE*^) strain. The compound was not detected in any previous study, however its ringed, anhydride form (hexylaconitic acid D) was naturally produced at 19.84 mg/L from *A. niger* AN27^24^. Compared to the hexylaconitic acid A titre in our study, this represents an increase of more than 400 fold. The increase of production of alkylcitric acids in this study is dramatic, given that SMs normally require years of research to reach similar titres^55–56^.

One reason for this steep increase from natural and artificial production may be due to the upregulation of genes involved in the production of building blocks of the fatty acids (acetyl-CoA and malonyl-CoA)^57^ and the citric acid and citric acid derived moieties (oxaloacetic acid). These building blocks can be generated from the breakdown of citric acid due to the action of a citrate lyase or similar enzyme^58^ which may be upregulated following the upregulation of *akcR*. Since *A. niger* is a well-known producer of citric acid, a high yield of the alkylcitric acids was expected. In this context, an even higher production of alkylcitric acids may be achieved if the *akcR* is overexpressed in an industrial citric acid producing strain of *A. niger*.

### Conclusion

Our study has shown that using a combination of bioinformatics and chemical structure, orphan compounds can be traced back to their biosynthetic gene cluster. Activation of these gene clusters may lead to the overproduction of orphan compounds as well as the production of novel derivatives. Moreover, modification of the expression level of tailoring genes may be able to shift the biosynthetic pathway toward desired products. This strategy was applied to locate the biosynthetic gene cluster of the alkylcitric acids. Overexpressing the co-localized transcription factor *akcR* and an unlinked tailoring gene (*cadA*), we overproduced alkylcitric acids and obtained six previously unreported alkylcitric acids.

## References

1. Fleming, A., On the antibacterial action of cultures of a *Penicillium*, with special reference to their use in the isolation of *B. influenzæ*. Br J Exp Pathol 1929, 10 (3), 226–236.

2. Rubinstein, A.; Lurie, Y.; Groskop, I., et al., Cholesterol-lowering effects of a 10 mg daily dose of lovastatin in patients with initial total cholesterol levels 200 to 240 mg/dl (5.18 to 6.21 mmol/liter). Am. J. Cardiol. 1991, 68 (11), 1123–1126.

3. Panda, D.; Rathinasamy, K.; Santra, M. K., et al., Kinetic suppression of microtubule dynamic instability by griseofulvin: implications for its possible use in the treatment of cancer. Proc. Natl. Acad. Sci. U.S.A. 2005, 102 (28), 9878–9883.

4. Gacek, A.; Strauss, J., The chromatin code of fungal secondary metabolite gene clusters. Appl. Microbiol. Biotechnol. 2012, 95 (6), 1389–1404.

5. Lind, A. L.; Wisecaver, J. H.; Lameiras, C., et al., Drivers of genetic diversity in secondary metabolic gene clusters within a fungal species. PLOS Biology 2017, 15 (11), e2003583.

6. Pi, B.; Yu, D.; Dai, F., et al., A genomics based discovery of secondary metabolite biosynthetic gene clusters in *Aspergillus ustus*. PLOS ONE 2015, 10 (2), e0116089.

7. Urlacher, V. B.; Girhard, M., Cytochrome P450 monooxygenases: an update on perspectives for synthetic application. Trends Biotechnol. 2012, 30 (1), 26–36.

8. Coleman, J. J.; Mylonakis, E., Efflux in fungi: la pièce de résistance. PLoS Pathog. 2009, 5 (6), e1000486.

9. Pitkin, J. W.; Panaccione, D. G.; Walton, J. D., A putative cyclic peptide efflux pump encoded by the *TOXA* gene of the plant-pathogenic fungus *Cochliobolus carbonum*. Microbiology 1996, 142, 1557–1565.

10. Desjardins, A. E.; Proctor, R. H., Molecular biology of *Fusarium* mycotoxins. Int. J. Food Microbiol. 2007, 119 (1-2), 47–50.

11. Fernandes, M.; Keller, N. P.; Adams, T. H., Sequence-specific binding by *Aspergillus nidulans* AflR, a C6 zinc cluster protein regulating mycotoxin biosynthesis. Mol. Microbiol. 1998, 28 (6), 1355–1365.

12. Zabala, A. O.; Xu, W.; Chooi, Y.-H., et al., Characterization of a silent azaphilone gene cluster from *Aspergillus niger* ATCC 1015 reveals a hydroxylation-mediated pyran-ring formation. Chem. Biol. 2012, 19 (8), 1049–1059.

13. Palys, S. A bioinformatics characterization of secondary metabolism and alkyl citric acid pathway reconstruction in *Aspergillus niger* NRRL3. M.S. Thesis, Concordia University, 2017.

14. Nielsen, K. F.; Mogensen, J. M.; Johansen, M., et al., Review of secondary metabolites and mycotoxins from the *Aspergillus niger* group. Anal Bioanal Chem 2009, 395 (5), 1225–1242.

15. Gil Girol, C.; Fisch, K. M.; Heinekamp, T., et al., Regio- and stereoselective oxidative phenol coupling in *Aspergillus niger*. Angew. Chem. Int. Ed. 2012, 51 (39), 9788–9791.

16. Fujii, R.; Matsu, Y.; Minami, A., et al., Biosynthetic study on antihypercholesterolemic agent phomoidride: General biogenesis of fungal dimeric anhydrides. Org. Lett. 2015, 17 (22), 5658–5661.

17. Fuchser, J.; Thiericke, R.; Zeeck, A., Biosynthesis of aspinonene, a branched pentaketide produced by *Aspergillus ochraceus*, related to aspyrone. J. Chem. Soc. Perkin Trans. 1995, 0 (13), 1663–1666.

18. Bode, Helge B.; Walker, M.; Zeeck, A., Cladospirones B to I from *Sphaeropsidales* sp. F-24′707 by variation of culture conditions. Eur. J. Org. Chem. 2000, 2000 (18), 3185–3193.

19. Clevenger, K. D.; Bok, J. W.; Ye, R., et al., A scalable platform to identify fungal secondary metabolites and their gene clusters. Nat. Chem. Biol. 2017, 13 (8), 895–901.

20. Oakley, C. E.; Ahuja, M.; Sun, W. W., et al., Discovery of McrA, a master regulator of *Aspergillus* secondary metabolism. Mol. Microbiol. 2017, 103 (2), 347–365.

21. Inglis, D. O.; Binkley, J.; Skrzypek, M. S., et al., Comprehensive annotation of secondary metabolite biosynthetic genes and gene clusters of *Aspergillus nidulans*, *A. fumigatus*, *A. niger* and *A. oryzae*. BMC Microbiol. 2013, 13, 91.

22. Hasegawa, Y.; Fukuda, T.; Hagimori, K., et al., Tensyuic acids, new antibiotics produced by *Aspergillus niger* FKI-2342. Chem. Pharm. Bull. 2007, 55 (9), 1338–1341.

23. Widenmuller, H. L.; Cavagna, F.; Fehlhaber, H. W., et al., 2-carboxymethyl-3-n-hexyl-maleic acid anhydride, a novel metabolite from an *Aspergillus*. Tetrahedron Lett. 1972.

24. Mondal, G.; Dureja, P.; Sen, B., Fungal metabolites from *Aspergillus niger* AN27 related to plant growth promotion. Indian J. Exp. Biol. 2000, 38 (1), 84–87.

25. Akira, I.; Washizu, M.; Kondo, K., et al., Isolation and identification of (+)-hexylitaconic acid as a plant growth regulator. Agric. Biol. Chem. 1984, 48 (10), 2607–2609.

26. Almassi, F.; Ghisalberti, E. L.; Rowland, C. Y., Alkylcitrate-derived metabolites from *Aspergillus niger*. J. Nat. Prod. 1994, 57 (6), 833–836.

27. Li, J. L.; Zhang, P.; Lee, Y. M., et al., Oxygenated hexylitaconates from a marine sponge-derived fungus *Penicillium* sp. Chem. Pharm. Bull. 2011, 59 (1), 120–123.

28. Koch, L.; Lodin, A.; Herold, I., et al., Sensitivity of *Neurospora crassa* to a marine-derived *Aspergillus tubingensis* anhydride exhibiting antifungal activity that is mediated by the MAS1 protein. Mar Drugs 2014, 12 (9), 4713–4731.

29. Storms, R.; Zheng, Y.; Li, H., et al., Plasmid vectors for protein production, gene expression and molecular manipulations in *Aspergillus niger*. Plasmid 2005, 53 (3), 191–204.

30. Master, E. R.; Zheng, Y.; Storms, R., et al., A xyloglucan-specific family 12 glycosyl hydrolase from *Aspergillus niger*: recombinant expression, purification and characterization. Biochem. J. 2008, 411 (1), 161–170.

31. Song, L.; Ouedraogo, J.-P.; Kolbusz, M., et al., Efficient genome editing using tRNA promoter-driven CRISPR/Cas9 gRNA in *Aspergillus niger*. PLOS ONE 2018, 13 (8), e0202868.

32. Sambrook, J.; Russell, D. W., Purification of nucleic acids by extraction with phenol:chloroform. CSH Protoc 2006, 2006 (1).

33. Aslanidis, C.; de Jong, P. J., Ligation-independent cloning of PCR products (LIC-PCR). Nucleic Acids Res 1990, 18 (20), 6069–6074.

34. Kelly, J. M.; Hynes, M. J., Transformation of *Aspergillus niger* by the *amdS* gene of *Aspergillus nidulans*. The EMBO J. 1985, 4 (2), 475–479.

35. Yolov, A. A.; Shabarova, Z. A., Constructing DNA by polymerase recombination. Nucleic Acids Res 1990, 18 (13), 3983–3986.

36. Toussaint, J.-P.; Pham, T. T. M.; Barriault, D., et al., Plant exudates promote PCB degradation by a rhodococcal rhizobacteria. Appl. Microbiol. Biotechnol. 2012, 95 (6), 1589–1603.

37. Ganzlin, M.; Rinas, U., In-depth analysis of the *Aspergillus niger* glucoamylase (*glaA*) promoter performance using high-throughput screening and controlled bioreactor cultivation techniques. J. Biotechnol. 2008, 135 (3), 266–271.

38. Klemke, C.; Kehraus, S.; Wright, A. D., et al., New secondary metabolites from the marine endophytic fungus *Apiospora montagnei*. J. Nat. Prod. 2003, 67 (6), 1058–1063.

39. Bentley, R.; Thiessen, C. P., Biosynthesis of itaconic acid in *Aspergillus terreus*. I. Tracer studies with C^14^-labeled substrates. J. Biol. Chem. 1957, 226 (2), 673–687.

40. Hossain, A. H.; Li, A.; Brickwedde, A., et al., Rewiring a secondary metabolite pathway towards itaconic acid production in *Aspergillus niger*. Microb Cell Fact. 2016, 15 (1), 130.

41. Stewart, M.; Capon, R. J.; Lacey, E., et al., Calbistrin E and two other new metabolites from an Australian isolate of *Penicillium striatisporum*. J. Nat. Prod. 2005, 68 (4), 581–584.

42. Wasil, Z.; Kuhnert, E.; Simpson, T. J., et al., Oryzines A & B, maleidride congeners from *Aspergillus oryzae* and their putative biosynthesis. J Fungi 2018, 4 (3).

43. Vanhanen, S.; West, M.; Kroon, J. T., et al., A consensus sequence for long-chain fatty-acid alcohol oxidases from *Candida* identifies a family of genes involved in lipid omega-oxidation in yeast with homologues in plants and bacteria. J. Biol. Chem. 2000, 275 (6), 4445–4452.

44. Schlömann, M.; Ngai, K. L.; Ornston, L. N., et al., Dienelactone hydrolase from *Pseudomonas cepacia*. J. Bacteriol. 1993, 175 (10), 2994–3001.

45. Walton, J. D., Horizontal gene transfer and the evolution of secondary metabolite gene clusters in fungi: an hypothesis. Fungal Genet. Biol. 2000, 30 (3), 167–171.

46. Robey, M. T.; Ye, R.; Bok, J. W., et al., Identification of the first diketomorpholine biosynthetic pathway using FAC-MS technology. Acs Chem Biol 2018, 13 (5), 1142–1147.

47. Harvey, C. J. B.; Tang, M.; Schlecht, U., et al., HEx: A heterologous expression platform for the discovery of fungal natural products. Sci Adv 2018, 4 (4), eaar5459.

48. Chiang, Y.-M.; Meyer, K. M.; Praseuth, M., et al., Characterization of a polyketide synthase in *Aspergillus niger* whose product is a precursor for both dihydroxynaphthalene (DHN) melanin and naphtho-γ-pyrone. Fungal Genet. Biol. 2011, 48 (4), 430–437.

49. Chiang, Y.-M.; Oakley, C. E.; Ahuja, M., et al., An efficient system for heterologous expression of secondary metabolite genes in *Aspergillus nidulans*. J. Am. Chem. Soc. 2013, 135 (20), 7720–7731.

50. Geib, E.; Brock, M., ATNT: an enhanced system for expression of polycistronic secondary metabolite gene clusters in *Aspergillus niger*. Fungal Biol Biotechnol 2017, 4, 13.

51. Chiang, Y.-M.; Szewczyk, E.; Davidson, A. D., et al., A gene cluster containing two fungal polyketide synthases encodes the biosynthetic pathway for a polyketide, asperfuranone, in *Aspergillus nidulans*. J. Am. Chem. Soc. 2009, 131 (8), 2965–2970.

52. Szewczyk, E.; Chiang, Y.-M.; Oakley, C. E., et al., Identification and characterization of the asperthecin gene cluster of *Aspergillus nidulans*. Appl. Environ. Microbiol. 2008, 74 (24), 7607–7612.

53. Bojja, R. S.; Cerny, R. L.; Proctor, R. H., et al., Determining the biosynthetic sequence in the early steps of the fumonisin pathway by use of three gene-disruption mutants of *Fusarium verticillioides*. J. Agric. Food Chem. 2004, 52 (10), 2855–2860.

54. Petersen, L. M.; Holm, D. K.; Knudsen, P. B., et al., Characterization of four new antifungal yanuthones from *Aspergillus niger*. J. Antibiot. 2015, 68 (3), 201–205.

55. Mulder, K. C. L.; Mulinari, F.; Franco, O. L., et al., Lovastatin production: From molecular basis to industrial process optimization. Biotechnol. Adv. 2015, 33 (6 Pt 1), 648–665.

56. Zhang, Y.-X.; Perry, K.; Vinci, V. A., et al., Genome shuffling leads to rapid phenotypic improvement in bacteria. Nature 2002, 415 (6872), 644.

57. Hopwood, D. A.; Sherman, D. H., Molecular genetics of polyketides and its comparison to fatty acid biosynthesis. Annu. Rev. Genet. 1990, 24, 37–66.

58. Chen, H.; He, X.; Geng, H., et al., Physiological characterization of ATP-citrate lyase in *Aspergillus niger*. J. Ind. Microbiol. Biotechnol. 2014, 41 (4), 721–731.

