## Supplementary material for "Biosynthesis of alkylcitric acids in *Aspergillus niger* involves both co-localized and unlinked genes"

Table S1. Primers used in this study.

| Primer Name | Description | Sequence 5' --> 3' |
| --- | --- | --- |
| PR_1 | NRRL3_11765 forward primer | TACTTCCAATCCAATCCATTTG ACGATATGGCTACTCCCCGCGCATC |
| PR_2 | NRRL3_11765 reverse primer | TTATCCACTTCCAATCCATTTG TCACTGCATCAATCCCAACGCAG |
| PR_3 | ANIp7 and ANIp9 reverse primer | AAATGGATTGGATTGGAAGTAC TGATCTAGCGTGTAATG |
| PR_4 | ANIp7 and ANIp9 forward primer | AAATGGATTGGAAGTGGATAAC TTAATTAAGTTTAAACG |
| PR_5 | NRRL3_00504 forward primer | TACTTCCAATCCAATCCATTTG ATGGTCACTATCACCGCCAAATC |
| PR_6 | NRRL3_00504 reverse primer | TTATCCACTTCCAATCCATTTG CTACTGCAACGGATTGACCAC |
| PR_7 | Cluster deletion 5' Flank forward primer | AAGCTTTCCTCTCAAGAGAC |
| PR_8 | Cluster deletion 5' Flank reverse primer | AAATGGATTGGAAGTGGATAAC CATACGAGTTATATCACTCTGC |
| PR_9 | Cluster deletion 3' Flank forward primer | TTATCCACTTCCAATCCATTTC GTCTGACGATTCGTCTGTG |
| PR_10 | Cluster deletion 3' Flank reverse primer | GTTGCACTACTTTGAACCG |
| PR_11 | CRISPR guide RNA NRRL3_11764 forward primer | TCTACTGCCACCATACCGGA GTTTTAGAGCTAGAAATAGCAAG |
| PR_12 | CRISPR guide RNA NRRL3_11764 reverse primer | TCCGGTATGGTGGCAGTAGA GACGAGCTTACTCGTTTCG |
| PR_13 | Screening of cluster deletion forward primer | CACGTACACAAACTCTACTGAG |
| PR_14 | Screening of cluster deletion reverse primer | CATTGCCTTTGCTTTGCCTTAC |
| PR_15 | NRRL3_11765 integration forward primer | GCAGTCGTCTGGGCAAGATTG |
| PR_16 | NRRL3_11765 integration Re | CACTTCCAGGAAGGGGATGAG |

*Overlapping sequences are underlined

Table S2: The NMR data for hexylaconitic acid A to C and hexylitaconic acid A to H

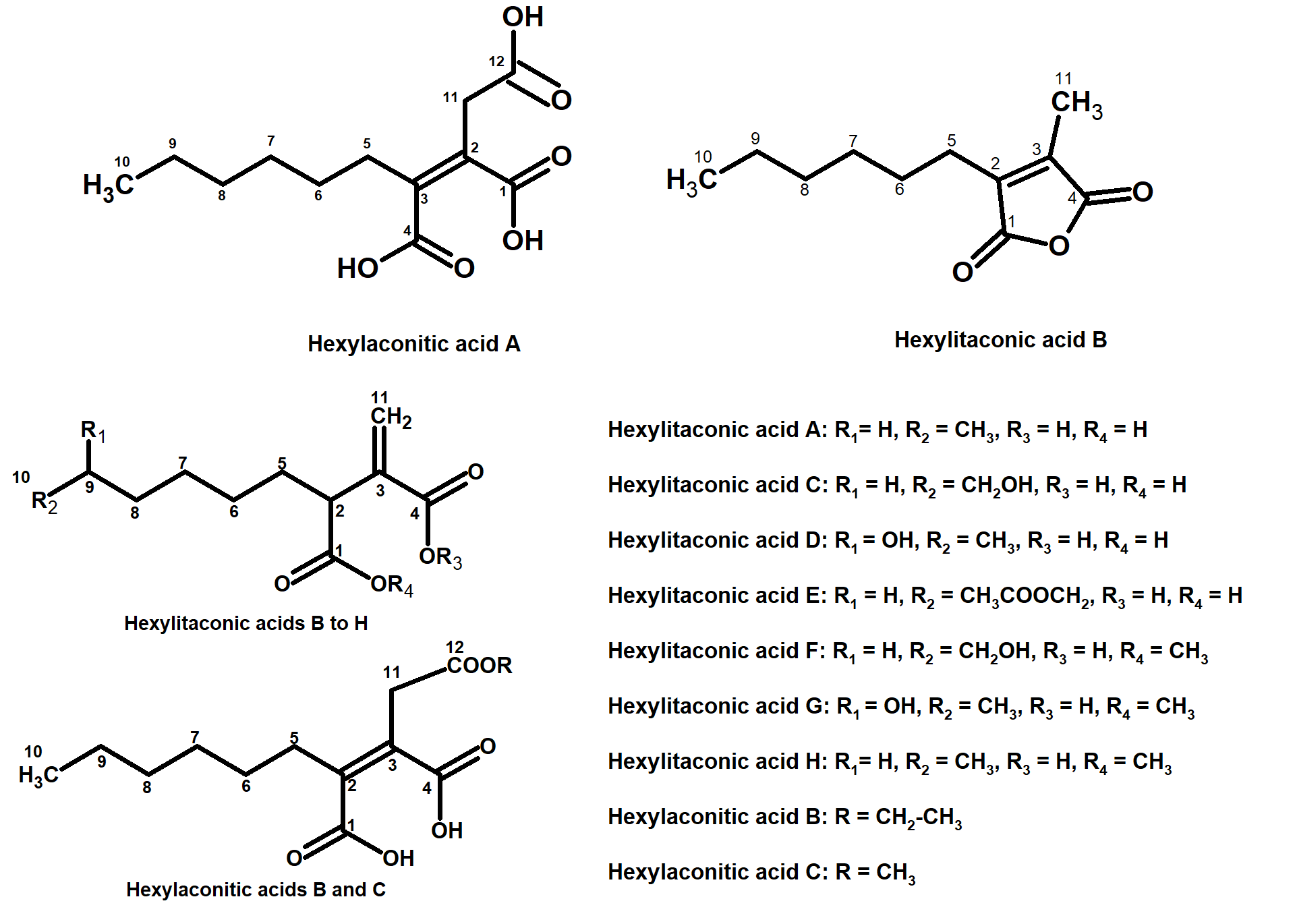

|  | **Hexylaconitic acid A (CDCl_3_)** | | | **Hexylaconitic acid B (CDCl_3_)** | | **Hexylaconitic acid C (CDCl_3_)** | |
| --- | --- | --- | --- | --- | --- | --- | --- |
|  | **^13^C** | | **^1^H** | **^13^C** | **^1^H** | **^13^C** | **^1^H** |
| 1 | 165.093 | |  | 165.314 |  | 165.231 |  |
| 2 | 147.950 | |  | 147.503 |  | 147.632 |  |
| 3 | 135.596 | |  | 136.355 |  | 136.158 |  |
| 4 | 165.07 | |  | 165.185 |  | 165.155 |  |
| 5 | 24.782 | | 2.491, t (8.0) | 24.877 | 2.483, t (7.5) | 24.892 | 2.476, t (8.0) |
| 6 | 27.309 | | 1.596, m | 27.495 | 1.592, m | 27.48 | 1.586, m |
| 7 | 28.971 | | 1.283 ÷ 1.303, m | 29.135 | 1.256 ÷ 1.323, m | 29.135 | 1.251 ÷ 1.317, m |
| 8 | 31.149 | | 1.283 ÷ 1.303, m | 31.297 | 1.256 ÷ 1.323, m | 31.29 | 1.251 ÷ 1.317, m |
| 9 | 22.24 | | 1.283 ÷ 1.303, m | 22.411 | 1.256 ÷ 1.323, m | 22.411 | 1.251 ÷ 1.317, m |
| 10 | 13.718 | | 0.882, t (7.0) | 13.957 | 0.889, t (7.0) | 13.949 | 0.885, t (7.0) |
| 11 | 29.1 | | 3.566, s | 29.582 | 3.504, s | 29.286 | 3.513, s |
| 12 | 172.56 | |  | 167.211 |  | 167.666 |  |
| O-CH_3_ |  | |  |  |  | 52.849 | 3.748, s |
| O-**CH_2_**-CH_3_ |  | |  | 62.054 | 4.205, q (7.0) |  |  |
| O-CH_2_-**CH_3_** |  | |  | 14.033 | 1.271, t (6.0) |  |  |
|  | **Hexylitaconic acid A (CDCl_3_)** | | | **Hexylitaconic acid B (CDCl_3_)** | | **Hexylitaconic acid C (CD_3_OD)** | |
|  | **^13^C** | | **^1^H** | **^13^C** | **^1^H** | **^13^C** | **^1^H** |
| 1 | 165.859 | |  | 179.571 |  | 175.709 |  |
| 2 | 140.415 | |  | 46.288 | 3.485, t (7.0) | 46.561 | 3.451, t (7.0) |
| 3 | 144.755 | |  | 136.909 |  | 139.428 |  |
| 4 | 166.253 | |  | 171.649 |  | 168.105 |  |
| 5 | 24.426 | | 2.451, dt (7.0; 0.5) | 30.565 | 5a: 1.904, m  5b: 1.706, m | 30.769 | 5a: 1.852, m |
|  |  |  |  |  |  |  | 5b: 1.684, m |
| 6 | 27.537 | | 1.574, m | 27.279 | 1.368 ÷ 1.245, m | 27.241 | 1.323 ÷ 1.369, m |
| 7 | 29.077 | | 1.304, m | 28.888 | 1.368 ÷ 1.245, m | 28.857 | 1.323 ÷ 1.369, m |
| 8 | 31.331 | | 1.304, m | 31.521 | 1.368 ÷ 1.245, m | 25.328 | 1.323 ÷ 1.369, m |
| 9 | 22.438 | | 1.304, m | 22.529 | 1.368 ÷ 1.245, m | 32.113 | 1.518, m |
| 10 | 13.969 | | 0.889, t (7.0) | 13.992 | 0.874, t (7.0) | 61.556 | 3.532, t (7.0) |
| 11 | 9.477 | | 2.069, s | 130.231 | 11a: 6.550, s  11b: 5.910, s | 125.625 | 11a: 6.329, s |
|  |  |  |  |  |  |  | 11b: 5.757, s |
|  | **Hexylitaconic acid D (CD_3_OD)** | | | **Hexylitaconic acid E (CDCl_3_)** | | **Hexylitaconic acid F (CD_3_OD)** | |
|  | **^13^C** | | **^1^H** | **^13^C** | **^1^H** | **^13^C** | **^1^H** |
| 1 | 175.703 | |  | 179.156 |  | 174.246 |  |
| 2 | 46.642 | | 3.453, t (8.0) | 46.877 | 3.148, t (7.0) | 46.596 | 3.529, t (7.0) |
| 3 | 139.49 | |  | 137.296 |  | 139.015 |  |
| 4 | 168.137 | |  | 171.461 |  | 167.864 |  |
| 5 | 30.812 | | 5a: 1.857, m | 29.688 | 5a: 1.928, m | 30.622 | 5a: 1.857, m |
|  |  |  | 5b: 1.691, m |  | 5b: 1.735, m |  | 5b: 1.679, m |
| 6 | 27.321 | | 1.318 ÷ 1.459, m | 27.192 | 1.349, m | 27.109 | 1.306 ÷ 1.364, m |
| 7 | 25.196 | | 1.318 ÷ 1.459, m | 28.884 | 1.349, m | 28.801 | 1.306 ÷ 1.364, m |
| 8 | 38.545 | | 1.318 ÷ 1.459, m | 25.666 | 1.349, m | 25.295 | 1.306 ÷ 1.364, m |
| 9 | 67.063 | | 3.695, m | 28.476 | 1.607, m | 32.11 | 1.514, m |
| 10 | 22.055 | | 1.133, d (7.0) | 64.551 | 4.043 t (7.0) | 61.501 | 3.529, t (7.0) |
| 11 | 125.587 | | 11a: 6.327, s | 129.609 | 11a: 6.529, s | 125.762 | 11a: 6.322, s |
|  |  |  | 11b: 5.756, s |  | 11b: 5.837, s |  | 11b: 5.741, s |
| O-CH_3_ |  | |  | 171.355  (CH_3_-**C**=O) |  | 50.96 | 3.648, s |
|  |  | |  | 20.992  (**CH_3_**-C=O) | 2.043, s |  |  |
|  | **Hexylitaconic acid G (CD_3_OD)** | | | **Hexylitaconic acid H (CDCl_3_)** | |  |  |
|  | **^13^C** | | **^1^H** | **^13^C** | **^1^H** |  |  |
| 1 | 174.246 | |  | 173.783 |  |  |  |
| 2 | 46.619 | | 3.477, t (7.0) | 46.202 | 3.488, t (7.0) |  |  |
| 3 | 139.315 | |  | 137.812 |  |  |  |
| 4 | 167.955 | |  | 171.142 |  |  |  |
| 5 | 30.683 | | 5a: 1.880, m | 31.214 | 5a: 1.886, m |  |  |
|  |  |  | 5b: 1.690, m |  | 5b: 1.676, m |  |  |
| 6 | 27.207 | | 1.125 ÷ 1.436, m | 27.389 | 1.298 ÷ 1.210, m |  |  |
| 7 | 25.151 | | 1.125 ÷ 1.436, m | 28.945 | 1.298 ÷ 1.210, m |  |  |
| 8 | 38.522 | | 1.125 ÷ 1.436, m | 31.57 | 1.298 ÷ 1.210, m |  |  |
| 9 | 67.002 | | 3.693, m | 22.54 | 1.298 ÷ 1.210, m |  |  |
| 10 | 22.039 | | 1.131, d (7.0) | 14.01 | 0.869, t (7.0) |  |  |
| 11 | 125.678 | | 11a: 6.315, s | 129.063 | 11a: 6.496, s |  |  |
|  |  |  | 11b: 5.735, s |  | 11b: 5.862, s |  |  |
| O-CH_3_ | 50.96 | | 3.649, s | 52.075 | 3.682, s |  |  |

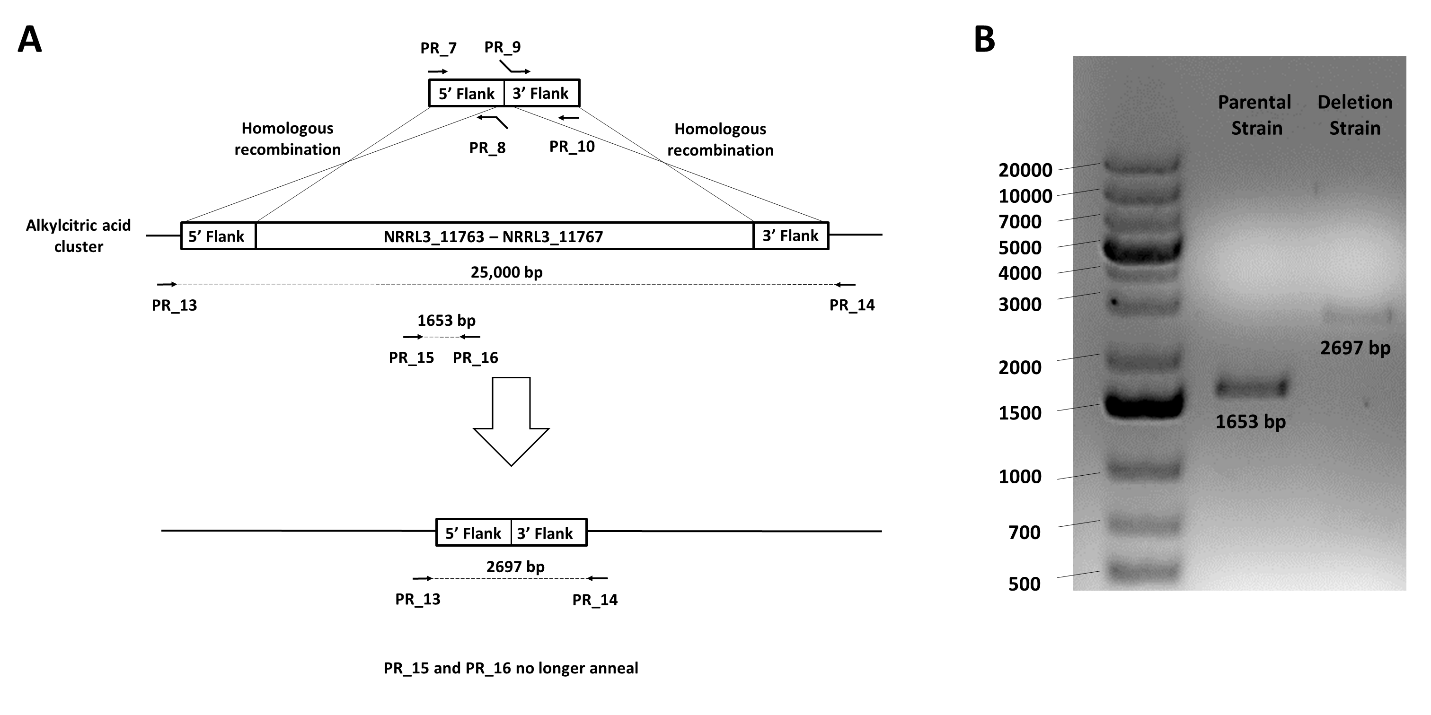

Figure S1. Diagram showing the deletion of genes NRRL_11763-7 by homologous recombination (A). Primers used are indicated by arrows and are listed in Table S1. Analysis by PCR confirming of the deletion of the alkylcitric acid cluster (ΔNRRL3_11763-7) in the SP1 (NRRL3_11765^OE^) strain (B).

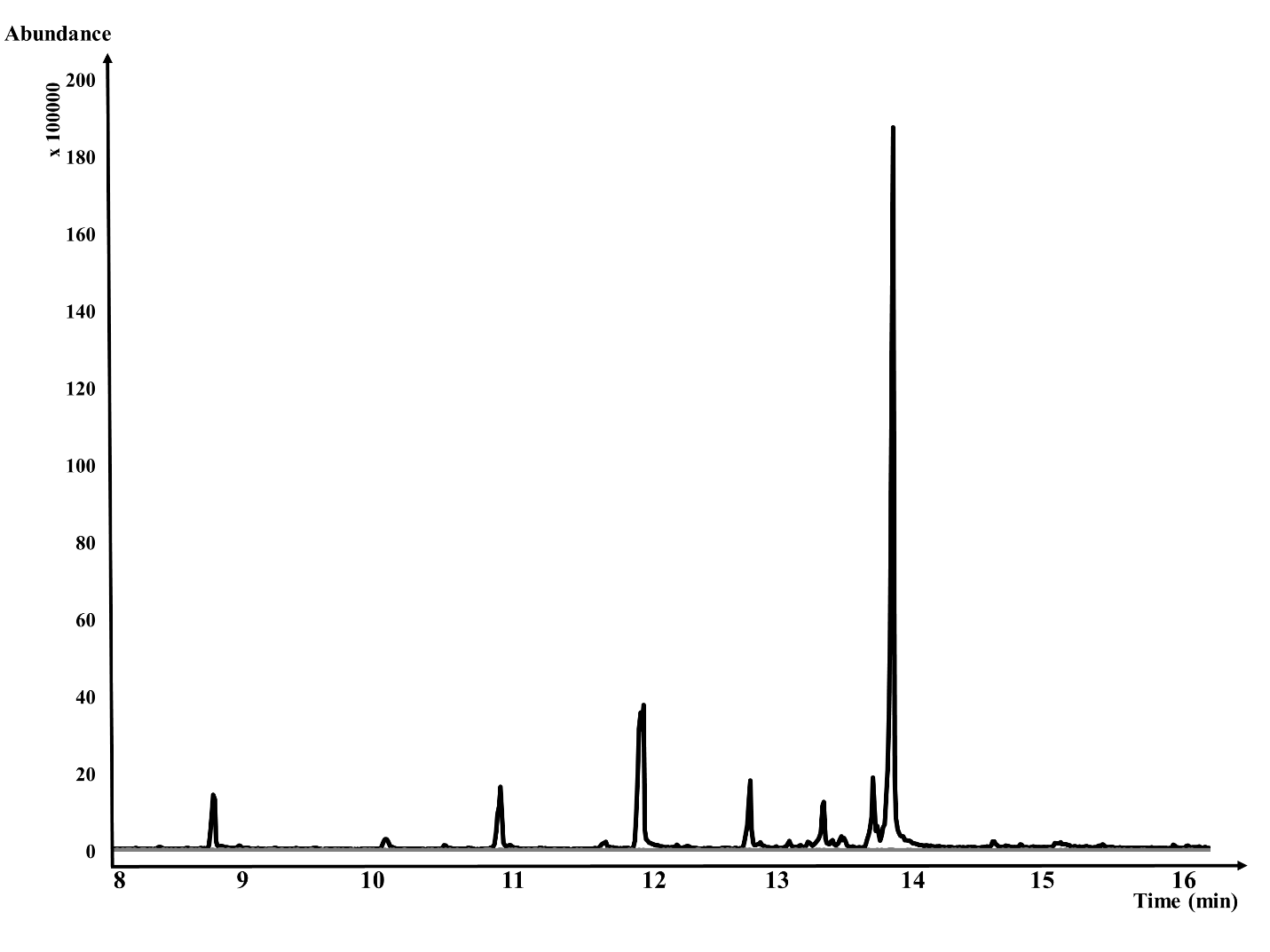

Figure S2. The GC-MS total ion chromatograms of strain SP1 (*akcR*^OE^) (in black) overlapping with parental strain *A. niger* PY 11 (in grey).

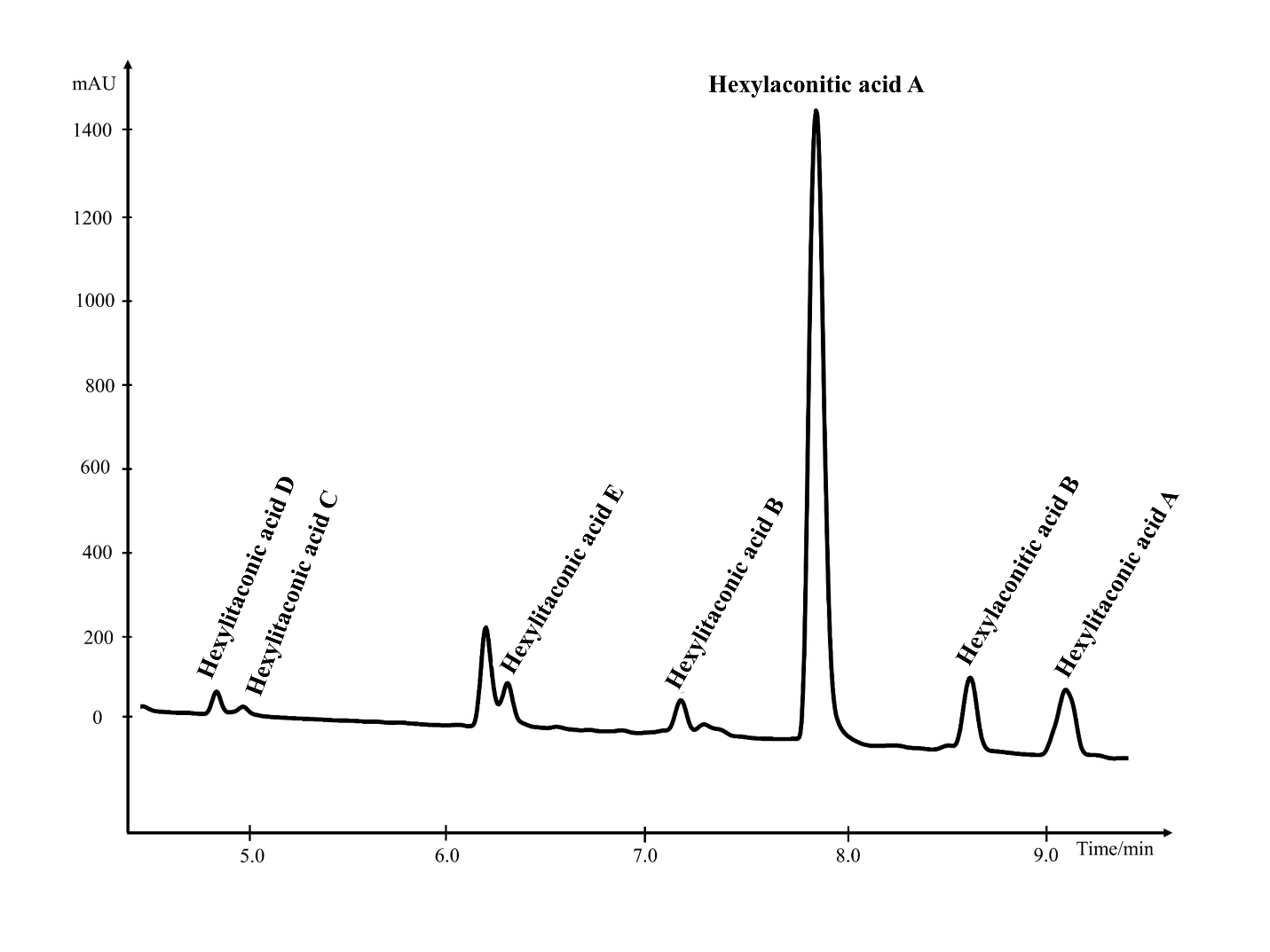

Figure S3. The HPLC total ion chromatograms of strain SP1 (*akcR*^OE^).

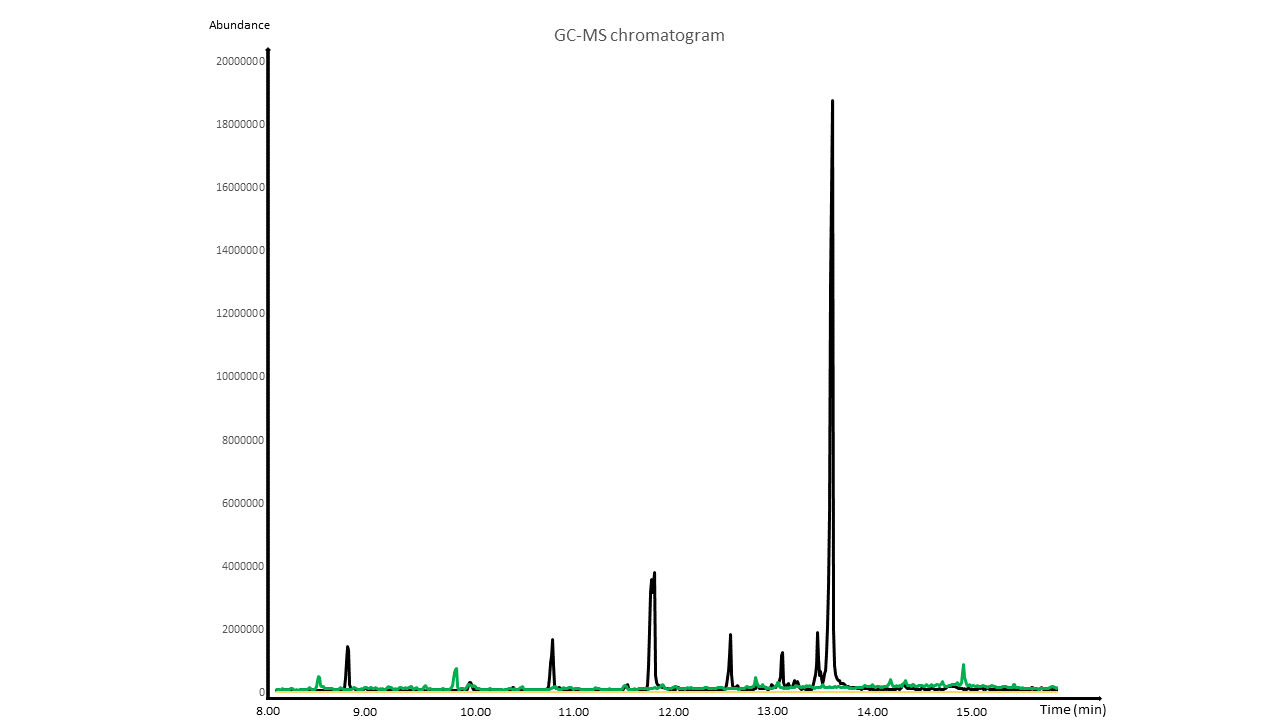

Figure S4. The GC-MS total ion chromatograms of the SP1 (*akcR^OE^*) (in black) overlapping with parental strain *A. niger* PY 11 (in yellow), and with the SP3 (*akcR^OE^* ΔNRRL3_11763-11767) strain (in green).

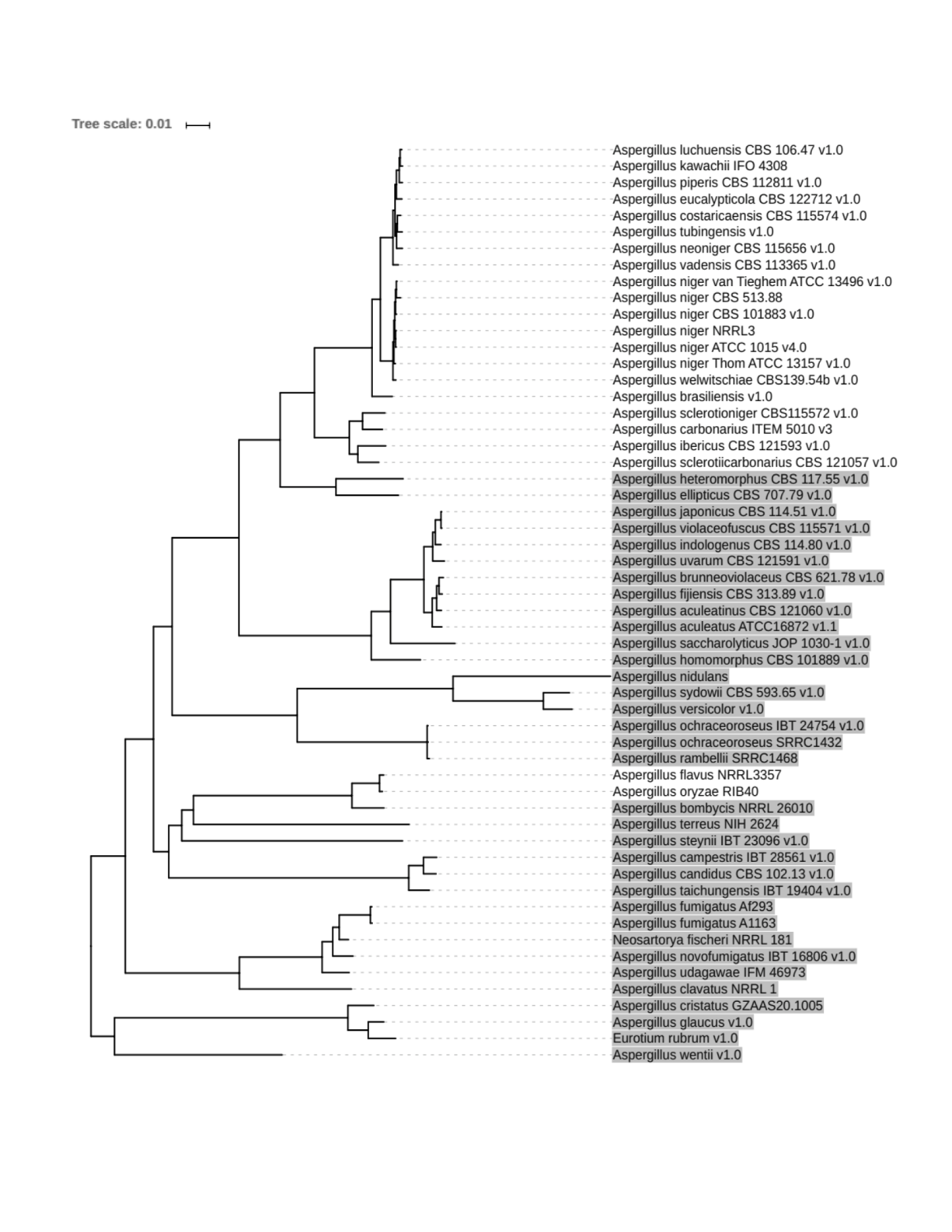

Figure S5: Taxonomic tree of published Eurotiomycetes genomes (<https://genome.jgi.doe.gov/programs/fungi/index.jsf>). Twenty orthologous alkylcitric acid gene clusters of section Nigri and two of section Flavi (*A. flavus* and *A. oryzae*) are displayed as non-highlighted names.

**Structural elucidation process of hexylaconitic acid A and hexylitaconic acid B**

Hexylaconitic acid A has an accurate mass (m/z) of 259.1176 ± 1.3 mmu ([M+H]^+^) which indicates its molecular formula is C_12_H_18_O_6_. According to ^13^C-NMR and DEPT 135 spectra, compound 1 contains 12 carbons including 5 quaternary, 6 methylene, and 1 methyl group. Of the 5 quaternary carbons, 3 are of a carbonyl (δC 173.319, 165.138, 165.115) and 2 are of an alkene (δC 148.102, 135.490). Since there is no ^13^C signal in the 40 - 130 ppm region, this suggests that there is no methylene or methyl group that is directly bonded to an oxygen atom. Therefore, the 3 carbonyl groups are defined as 3 carboxylic acid groups which is consistent with the presence of 6 oxygen atoms in the molecular formula. Five methylene groups (C5-C9) were shown by COSY-NMR and HMBC-NMR spectra to be chained together forming a saturated hydrocarbon tail (-C_6_H_13_) which ends at a methyl group (C10). The -C_6_H_13_ structural element is also confirmed by the splitting pattern of these protons. This partial structure was extended by observing the correlations between the saturated hydrocarbon chain, the carbonyl group (C4) and 2 alkene groups (C2, C3) on HMBC. According to ^1^H-NMR, the last methylene group (C11) has a singlet signal (δH 3.549, 2H) which indicates it is separated from the other 5 methylene groups. The HMBC-NMR spectrum determined that this -CH_2_- group has correlations with 2 carbonyl groups (C1, C12) and 2 alkene groups (C2, C3) but has no correlation with the –C_6_H_13_ chain. On the basis of the previously mentioned NMR evidences, the structure was determined to be hexylaconitic acid A. The ^1^H and ^13^C-NMR chemical shifts are listed in Table S2. The selected COSY and HMBC correlations are shown in Figure S6.

Hexylitaconic acid A has molecular formula of C_12_H_16_O_3_ and an accurate mass of m/z 209.1183 ± 1.0 mmu ([M+H]^+^). The ^13^C-NMR and DEPT 135 spectra indicate that compound 2 contains 12 carbons including 4 quaternary, 5 methylene, and 2 methyl groups. The ^1^H and ^13^C-NMR chemical shifts reveal that compound 2 also contains a saturated hydrocarbon chain (-C_6_H_13_) and it is identical to that of compound 1. The rest of the structure consists of 4 quaternary carbons and 1 methyl group. Three of these 4 quaternary carbons (C2, C3, C4; δC 144.755, 140.415, 165.859, respectively) have HMBC correlations with the methylene group (C5, δC 24.426) of the hydrocarbon chain. The methyl group has a singlet signal (δH 2.069, 3H) on the ^1^H-NMR spectrum which is evidence that there is no correlation between itself and nearby protons. Moreover, according to the HMBC spectrum, this methyl group has correlations with 3 quaternary carbons (C2, C3, C1; δC 144.755, 140.415, 166.253, respectively). On the basis of the NMR evidences and its molecular formula, the only structure that is suitable was hexylitaconic acid A. Selected HMBC correlations are shown in Figure S6 and ^1^H and ^13^C NMR chemical shifts are listed in Table S2.

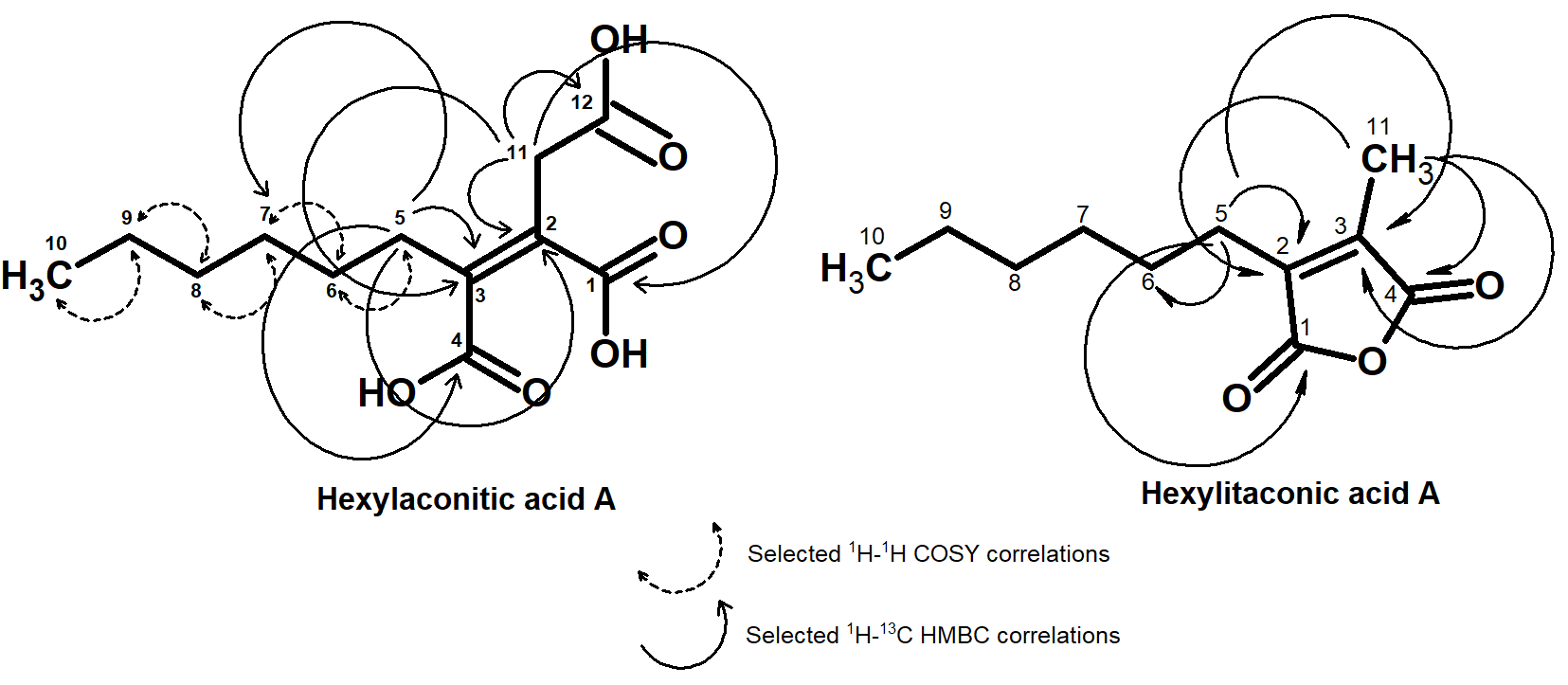

Figure S6. Structure and selected HMBC and COSY correlations of hexylaconitic acid A and hexylitaconic acid A

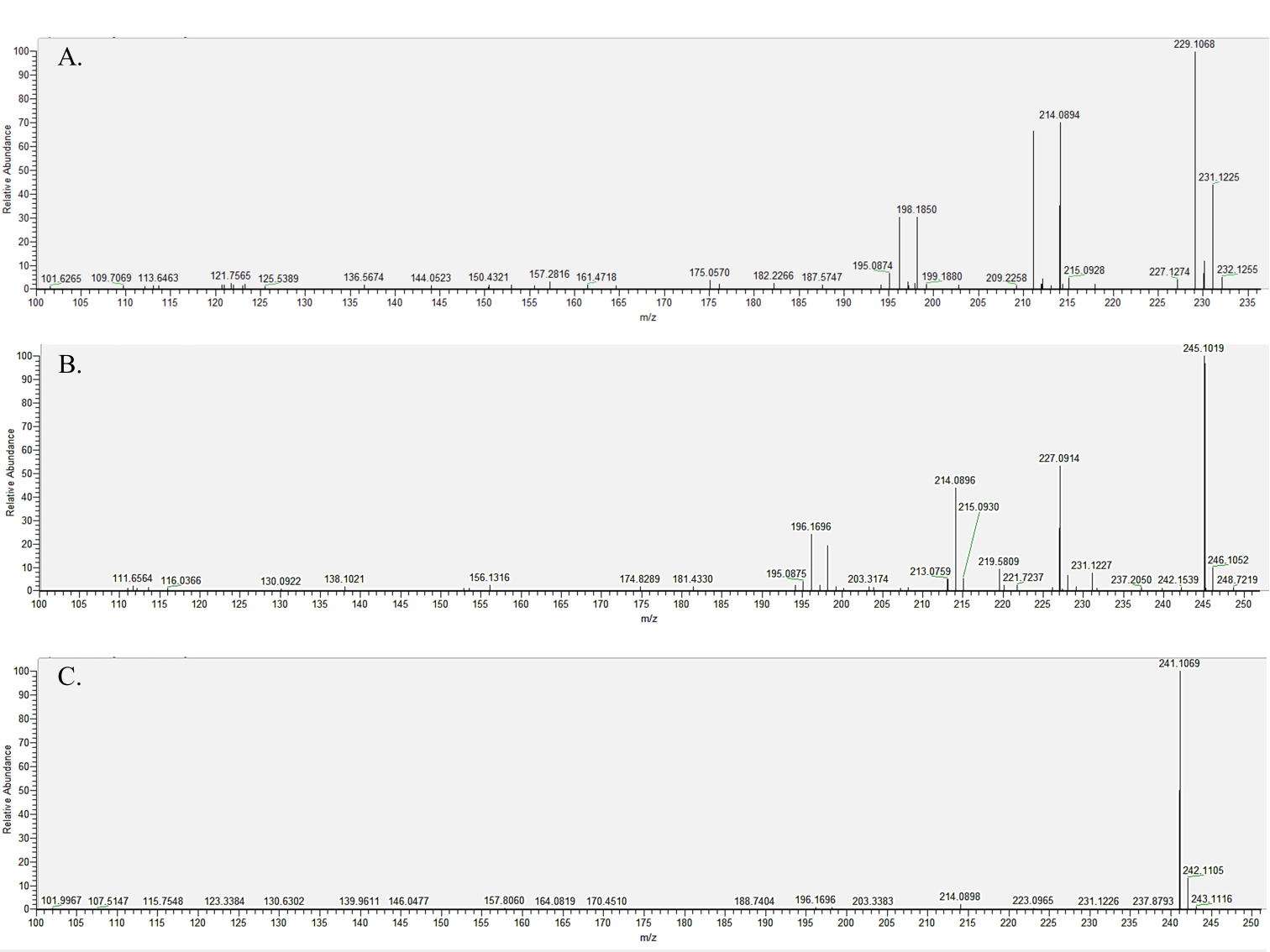

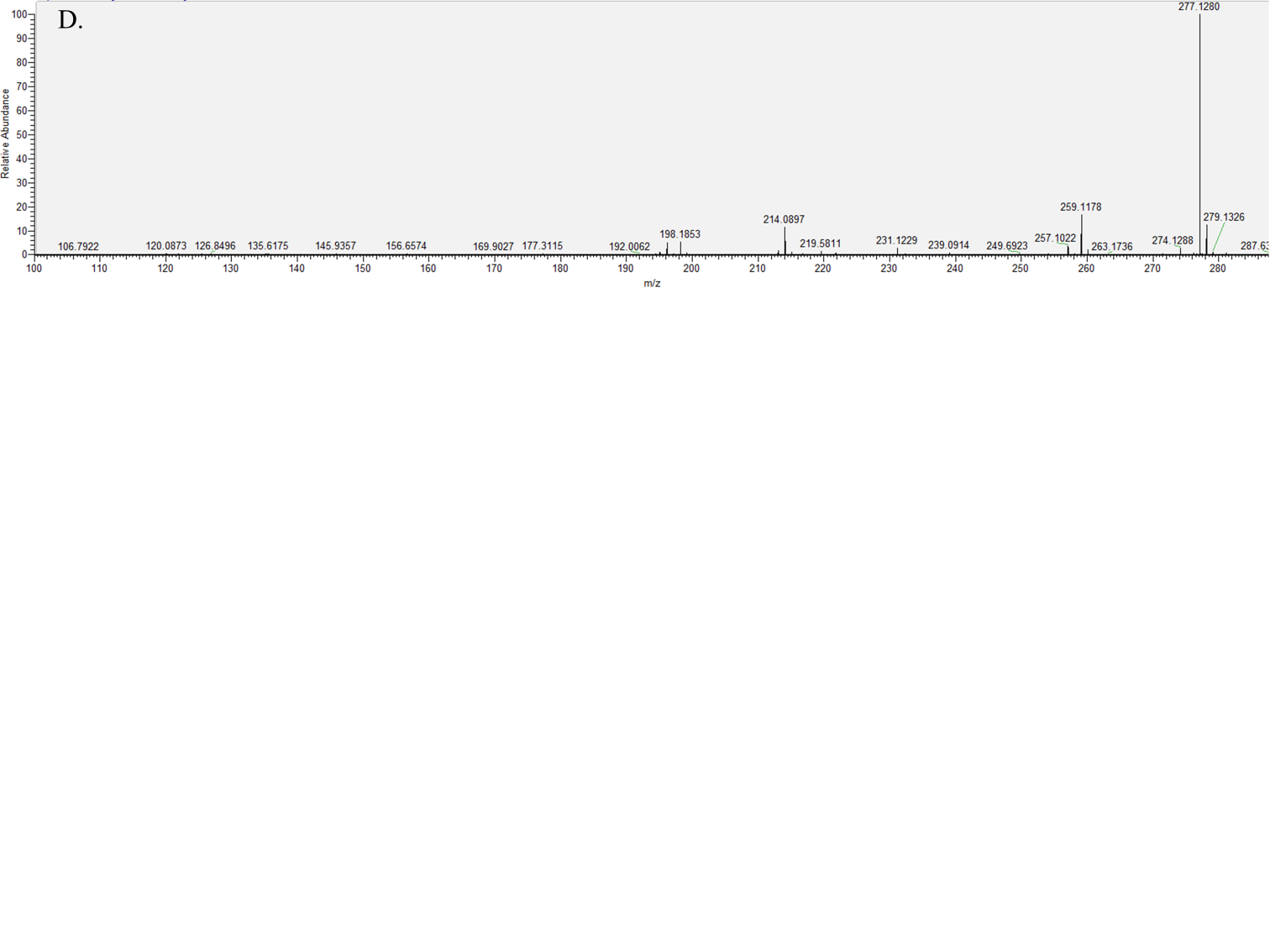

Figure S7. Mass spectra (LC-MS) of hexylitaconic acid I ([M+H] ^+^ = 229.1076 ± 1.1 mmu) (A), hexylitaconic acid J ([M+H] ^+^ = 245.1025 ± 1.2 mmu) (B), hexylaconitic acid D ([M+H] ^+^ = 241.1076 ± 1.2 mmu) (C), and hexylcitric acid ([M+H]^+^ = 277.1287 ± 1.4 mmu) (D).

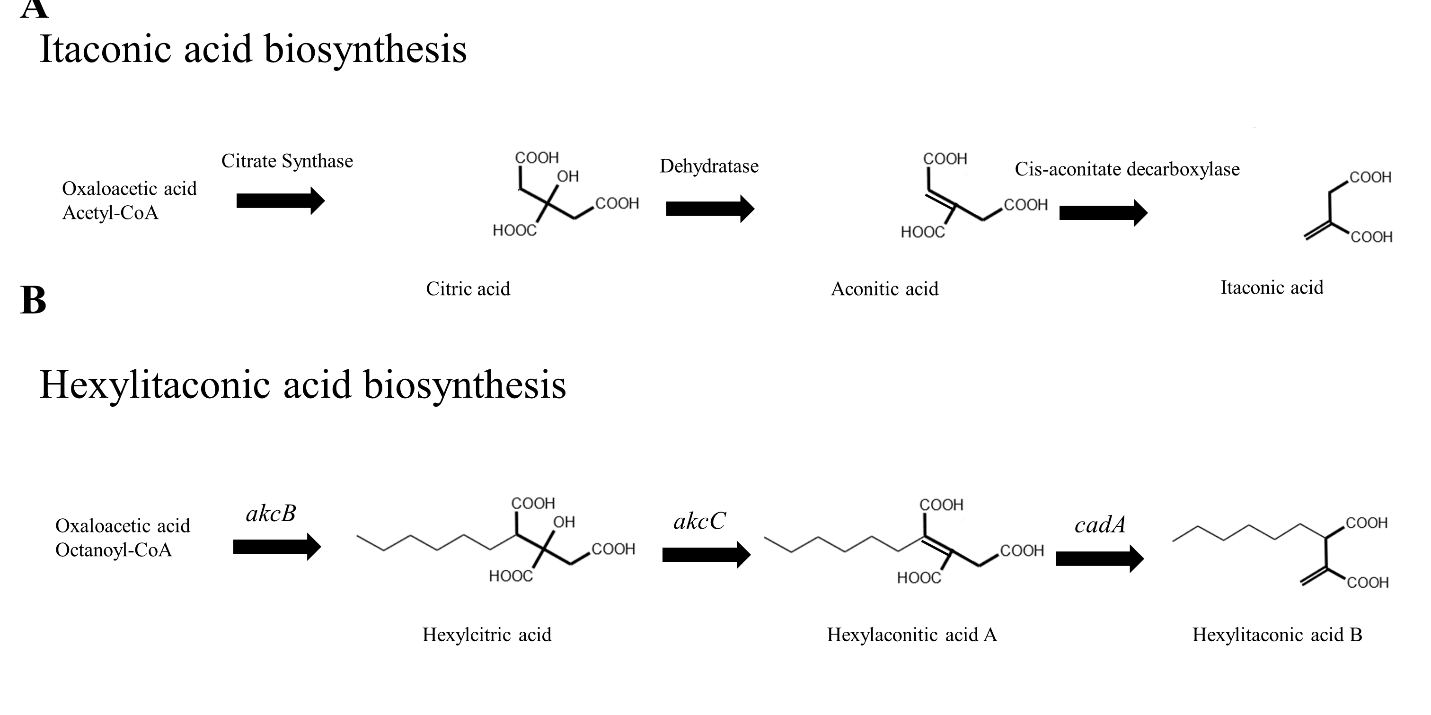

Figure S8. Biosynthetic pathways of itaconic acid (Bentley, 1957; Hossain, 2016) (A), the predicted hexylitaconic acid B pathway (B). Thick bonds indicate the common moieties between the two pathways.
